# Control of noncoding RNA production and histone levels by a 5’ tRNA fragment

**DOI:** 10.1101/498949

**Authors:** Ana Boskovic, Xin Yang Bing, Ebru Kaymak, Oliver J. Rando

## Abstract

Small RNAs derived from mature tRNAs, referred to as tRNA fragments or “tRFs”, are an emerging class of regulatory RNAs with poorly understood functions in cellular regulation. We recently identified a role for one specific tRF - 5’ tRF-Gly-GCC, or tRFGG - in repression of genes associated with the endogenous retroelement MERVL, but the mechanistic basis for this regulation was unknown. Here, we show that tRF-GG plays a role in production of a wide variety of noncoding RNAs normally synthesized in Cajal bodies. Among these noncoding RNAs, tRF-GG regulation of the U7 snRNA modulates heterochromatin-mediated transcriptional repression of MERVL elements by supporting an adequate supply of histone proteins. Importantly, the effects of inhibiting tRF-GG on histone mRNA levels, activity of a histone 3’ UTR reporter, and ultimately on MERVL regulation could all be suppressed by the U7 RNA. We show that the related RNA-binding proteins hnRNPF and H bind directly to tRF-GG, and are required for Cajal body biogenesis. Together, our data reveal a conserved mechanism for 5’ tRNA fragment control of noncoding RNA biogenesis and, consequently, in global chromatin organization.

## INTRODUCTION

It has been known for some time that mature tRNAs can be cleaved in response to cellular stressors (Lee and Collins, 2005), but only recently have the resulting cleavage products - broadly known as tRNA fragments, or tRFs - been appreciated as potential regulatory molecules in their own right (Keam and Hutvagner, 2015). Although tRNA fragments have in some cases been reported to function in complex with Argonaute proteins and act effectively like microRNAs or endo-siRNAs (Deng et al., 2015; Martinez et al., 2017; Schorn et al., 2017), they have also been reported to have Argonaute-independent regulatory functions ranging from inhibition of translation to control of apoptosis (Couvillion et al., 2012; Elbarbary et al., 2009; Gebetsberger et al., 2012; Goodarzi et al., 2015; Ivanov et al., 2011; Kim et al., 2017; Molla-Herman et al., 2015; Sobala and Hutvagner, 2013; Zhang et al., 2009). The diversity of proposed mechanisms for tRF function in part reflects the multitude of types of tRNA fragments that have been identified - 22 nt fragments derived from the 3’ ends of mature tRNAs have been found associated with Argonaute proteins (Kumar et al., 2014; Kuscu et al., 2018) and have been suggested to direct cleavage of retrotransposon RNAs (Martinez et al., 2017; Schorn et al., 2017), whereas longer (28-32 nt) fragments arising from tRNA 5’ ends appear to play more diverse mechanistic roles. For instance, 5’ fragments of valine tRNAs serve as global repressors of translation in archaea, yeast, and mammals, and in some cases appear to act by interfering with translational initiation (Bakowska-Zywicka et al., 2016; Gebetsberger et al., 2017; Guzzi et al., 2018; Luo et al., 2018).

We previously showed that interfering with a 5’ fragment of tRNA-Gly-GCC (hereafter, tRF-GG) using an antisense LNA oligonucleotide resulted, in both murine ES cell culture and in preimplantation embryos, in derepression of ~50 genes associated with the long terminal repeat (LTR) of the endogenous retroelement MERVL (Sharma et al., 2016). This functional link between a tRNA fragment and LTR element control is particularly interesting given the ancient and widespread role for tRNAs in LTR element replication - tRNAs almost universally serve as primers for reverse transcription of LTR elements (Marquet et al., 1995) - as well as recent studies reporting that 3’ tRNA fragments can interfere with multiple stages of the LTR element life cycle (Deng et al., 2015; Martinez et al., 2017; Schorn et al., 2017). In the case of tRF-GG-mediated control of MERVL elements, however, we find no identifiable homology between the 5’ tRF-GG and either the LTR or the primer binding sequence of MERVL (which is primed by homology to <u>L</u>eucine tRNAs), making it unlikely that MERVL regulation occurs through homology-directed RNA targeting.

Here, we set out to uncover the mechanistic basis for repression of MERVL-associated genes by tRF-Gly-GCC. To our surprise, we find that control of MERVL elements is a downstream result of an evolutionarily-conserved function for tRF-GG in supporting noncoding RNA production. Manipulation of tRF-GG levels in human and mouse ES cells affects the levels of a wide range of noncoding RNAs, including snoRNAs, scaRNAs, and various U RNAs, all of which are normally produced in a subnuclear organelle known as the Cajal body. One such RNA, the U7 noncoding RNA, is essential for 3’ UTR processing of histone pre-mRNAs. We show that tRF-GG control of U7 levels has downstream effects on histone mRNA and protein levels, as well as global chromatin compaction, and that the effects of tRF-GG on histones and on MERVL target gene transcription can be suppressed by manipulating U7 snRNA levels. Finally, we identify the related proteins hnRNP F/H as direct binding partners for tRF-GG, and show that these RNA-binding proteins are required for normal Cajal body biogenesis and for repression of MERVL-driven gene expression. Taken together, our data reveal a novel pathway for tRNA fragment function in mammals, linking tRNA cleavage to regulation of noncoding RNA production.

## RESULTS

### Chromatin-mediated repression of MERVL target transcription by tRF-GG

We recently identified a role for a 28 nt RNA derived from the 5’ end of tRNA-Gly-GCC - tRF-Gly-GCC, hereafter referred to as tRF-GG for brevity - as a repressor of genes associated with the long terminal repeat (LTR) of the endogenous retroelement MERVL (Sharma et al., 2016). This repressive activity does not appear to be a consequence of sequence homology between tRF-GG and the MERVL element, based on three observations: 1) there is no significant sequence homology between tRF-GG and MERVL (which uses tRNA-Leu for replication), and many of the tRF-GG target genes are in any event regulated by “solo” LTRs which have lost the MERVL primer binding sequence; 2) LNAs targeting the 3’ end of tRNA-Gly-GCC have no effect on MERVL target expression (Sharma et al., 2016); 3) transfection of ES cells with various synthetic 3’ tRNA fragments of potential relevance to either tRF-Gly-GCC or to MERVL has no significant effect on MERVL target gene expression (**Supplemental Figure S1A**), arguing against tRF-GG affecting levels or function of other tRNA fragments that might have more direct roles in MERVL control. Taken together, these data do not support a role for Argonaute-mediated targeting of MERVL mRNA (or DNA) sequences in tRF-GG regulation of this gene set.

Based on our initial studies suggesting that tRF-GG mediated repression of MERVL occurs at the level of transcription rather than RNA stability (Sharma et al., 2016), we set out to further distinguish between tRF-GG regulation of target gene synthesis vs. RNA decay. Metabolic labeling with 4-thiouridine was used to label newly-synthesized RNAs in murine E14 ESC cultures following inhibition of tRF-GG function using an LNA-containing antisense oligo (**Supplemental Table S1**). Inhibition of tRF-GG (“tRF-GG KD”) resulted in dramatically increased levels of total MERVL target mRNA abundance, replicating our prior findings (Sharma et al., 2016). Importantly, we also observed increased levels of MERVL target genes following tRF-GG inhibition in the purified 4SU-labeled, newly-synthesized, mRNA fraction, indicating that tRF-GG represses target gene synthesis rather than stability (**Figure 1A**, **Supplemental Figure S1B**). In addition, as previously reported, MERVL target derepression was accompanied by increased RNA Pol2 levels on target genes and could be recapitulated using LTR-driven tdTomato fluorescent reporter (Sharma et al., 2016). We conclude that tRF-Gly-GCC plays a role in *transcriptional* repression of MERVL LTRs.

**Figure 1.**
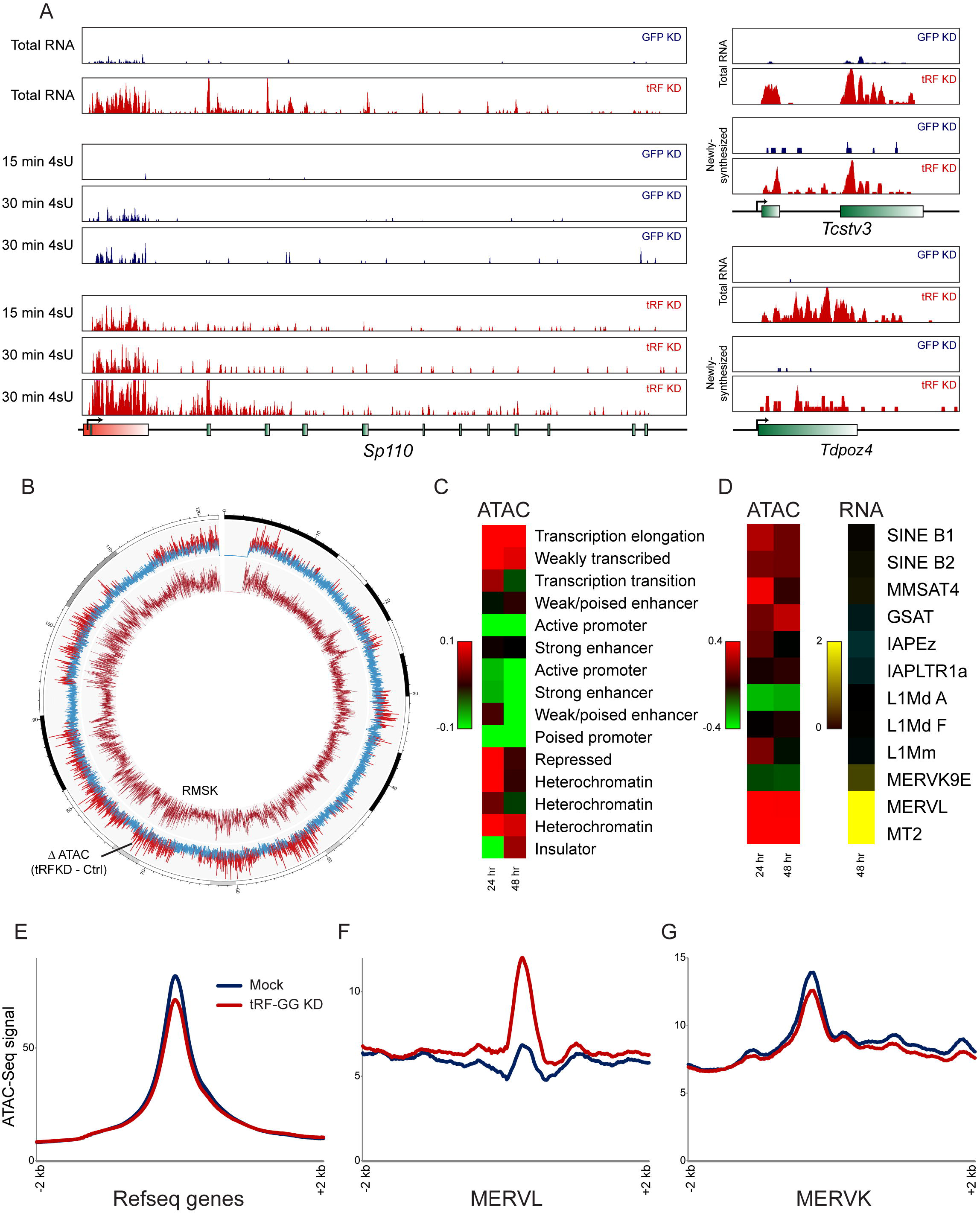
Transcriptional derepression of MERVL elements in response to tRF-Gly-GCC inhibition. A) Metabolic labeling reveals transcriptional derepression upon tRF-Gly-GCC inhibition. Genome browser tracks show total RNA levels, and newly-synthesized RNAs obtained after 15 or 30 minutes of 4-thiouridine (4SU) labeling, for ES cells transfected with esiRNAs targeting GFP, or an LNA oligonucleotide antisense to tRF-Gly-GCC. Effects of tRF inhibition on previously-described MERVL-associated target genes (Sharma et al., 2016) are nearly identical for total RNA as well as newly-synthesized RNA (see also **Supplemental Figure S1B**). B) Circos plot showing ATAC-Seq data for control and tRF-GG-inhibited ES cells across chromosome 11. Inner circle shows Repeatmasker density, and blue/red trace in the outer ring shows the change in ATAC signal between tRF KD and control ES cells, with red indicating more than 1.5 fold increased chromatin accessibility following tRF-GG inhibition. C) Increased accessibility at heterochromatin and weakly-transcribed regions in tRF-GG-inhibited ES cells. Heatmap shows log2 fold change in ATAC-Seq reads following tRF-GG inhibition, aggregated across the indicated types of chromatin (Bogu et al., 2015). D) As in (C), with tRF-GG effects on ATAC-Seq occupancy and RNA abundance averaged across the indicated repeat elements. E-G) Examples showing average ATAC-Seq signal across the indicated genomic elements - Refseq genes (E), MERVL elements (F), or MERVK elements (G).

Next, we asked how transcription of MERVL-driven genes is controlled. Like many retroelements, MERVL LTRs are packaged into, and repressed by, heterochromatin (Macfarlan et al., 2011). Interfering with chromatin assembly, for instance via knockdown of the CAF-1 histone chaperone (Ishiuchi et al., 2015), results in derepression of MERVL-driven transcripts. To investigate the effects of tRF-Gly-GCC on chromatin architecture, we carried out ATAC-Seq (Buenrostro et al., 2015) in mouse ES cells to measure changes in chromatin accessibility genome-wide upon tRF-GG KD (**Figure 1B**). Consistent with the enhanced transcription observed at MERVL LTRs, we find that inhibition of tRF-GG resulted in a broad increase in chromatin accessibility over MERVL elements and throughout heterochromatin (**Figures 1B-D**), with minimal changes in ATAC-Seq signal over euchromatic transcriptional start sites (**Figures 1E-G**). To extend these findings to a more developmentally-relevant scenario, we microinjected IVF-derived zygotes with a synthetic tRF-GG oligonucleotide to mimic the process of sperm delivery of tRFs to the zygote. We assayed chromatin accessibility in control and injected embryos by ATAC-qPCR, confirming that increasing tRF-GG levels resulted in a decrease in accessibility at MERVL elements (**Supplemental Figure S1C**). Thus, tRF-Gly-GCC manipulation alters chromatin accessibility in both ES cells and in preimplantation embryos.

### tRF-GG is a positive regulator of histone genes

To explore the mechanistic basis for global heterochromatin opening, we turned to RNA-Seq and ribosome footprinting data (Sharma et al., 2016) to identify potential effects of tRF-GG inhibition on expression of key chromatin regulators such as CAF-1. In addition, given the species-specific genomic locations of many ERVs such as MERVL, we also gathered RNA-Seq data in H9 human ES cells subject to tRF-GG inhibition, to identify conserved and divergent transcriptional consequences of tRF-GG inhibition. In both mouse and human ESCs, tRF-GG inhibition did not affect expression of any known chromatin regulators of MERVL such as CAF-1, Kdm1a, or Ehmt1 (**Supplemental Tables S1-2**). Intriguingly, tRF-GG inhibition in human ESCs had minimal effects on HERV expression, indicating that ERV regulation by this tRNA fragment is confined to mouse ES cells.

Instead, we identified two conserved molecular phenotypes resulting from tRF-GG inhibition in both human and mouse ES cells: repression of histone mRNAs (**Figures 2A-C**, **Supplemental Figure S2**), and decreased expression of a variety of noncoding RNAs (see below). Given our finding of global heterochromatin decompaction in tRF-inhibited cells (**Figures 1B-C**), and the common derepression of ERV elements in undercompacted genomes (Ishiuchi et al., 2015; Lenstra et al., 2011), we focus first on tRF-GG effects on histone genes. We confirmed by qRT-PCR that tRF-GG inhibition causes a decrease in histone mRNA abundance (**Figure 2D**), and that reduced histone mRNA levels are accompanied by a decrease in histone protein levels (**Supplemental Figures S2D-E**). Importantly, we find that direct tRF “overexpression” via transfection of ES cells with a synthetic 28 nt tRF-Gly-GCC also resulted in *increased* histone mRNA abundance (**Figure 2D**), demonstrating that repression of histone genes observed in response to tRF-GG inhibition does not represent an off-target gain of function exhibited by our anti-tRF-GG LNA oligo. Together, our gain and loss of function studies demonstrate that tRF-Gly-GCC plays a conserved role in histone mRNA expression, with the MERVL LTR representing a sensitized reporter for chromatin assembly specifically in murine ES cells.

**Figure 2.**
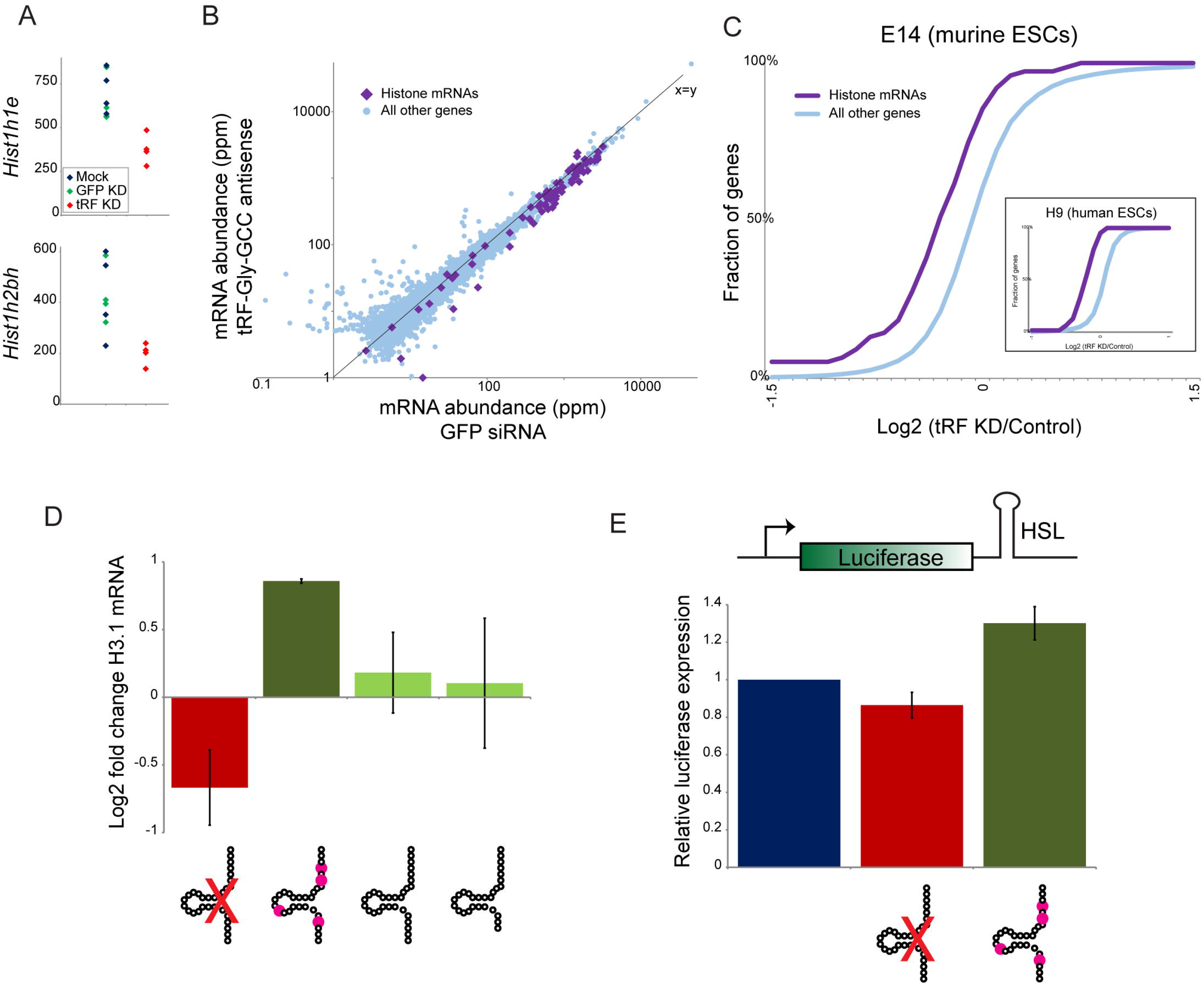
tRF-Gly-GCC represses expression of histone genes via the histone 3’ UTR. A) mRNA abundance for two example histone genes - *Hist1h1e* (top) or *Hist1h2bh* (bottom) - in four replicate RNA-Seq libraries from mock-transfected, GFP KD, and tRF-GG KD mES cells, as indicated. B) Scatterplot comparing RNA abundance for histone genes (purple diamonds) and all other genes in GFP KD ES cells (x axis) and tRF-GG-inhibited ES cells (y axis). Note that nearly all histone genes fall below the x=y diagonal. C) Cumulative distribution of the effects of tRF-GG inhibition on histone mRNA expression, with y axis showing cumulative fraction of genes exhibiting any given log2 fold change in expression (x axis). Main panel shows data from murine ES cells (n=4 replicates, KS p = 7.7e-5), while inset shows data for human ESCs. See also **Supplemental Figures S2A-C**. D) q-RT-PCR for *Hist2h3b* showing effects of transfecting the anti-tRF-GG LNA, a synthetic tRF-GG oligonucleotide bearing most of the expected nucleotide modifications present in tRNA-Gly-GCC, and two synthetic nucleotides without any modified nucleotides. Although the modified synthetic tRF was more active in this assay, we found no significant effect on tRF modification in several other assays (not shown) and so this was not pursued in further detail. E) tRF-GG regulates luciferase activity through a histone 3’ UTR. We generated stable ES cell lines carrying a luciferase reporter bearing the 3’ UTR of *Hist2h3b* (**Supplemental Figure S2F** shows data for an independent cell line bearing the *Hist1h4j* 3’ UTR). Bar graph shows average changes to reporter activity in response to control KD, tRF-GG LNA (14% decrease, p = 0.038), or the modified tRF-GG oligo (30% increase, p = 0.0002).

What is the mechanistic basis for tRF-GG-mediated repression of the histone genes? Histone expression is largely confined to the S phase of the cell cycle and could thus report on changes in cell cycle profile. However, FACS analysis of tRF-GG-inhibited ES cells revealed no change in the fraction of cells in S phase (**Supplemental Figure S3**). Histone expression is highly regulated at levels from transcription to translation; perhaps the most unique feature of histone expression is the role of several cis-acting RNA elements in the histone 3’ UTR - a short stem loop known as histone stem loop (HSL) that binds to stem loop binding protein (SLBP), and the histone downstream element (HDE) that binds to the U7 noncoding RNA and the associated U7 snRNP - in regulation of histone pre-mRNA processing (Dominski and Marzluff, 1999; Marzluff and Koreski, 2017). To separate the effects of tRF-GG manipulation on the histone 3’ UTR from effects on the histone promoter or coding sequence, we generated stable ES cell lines carrying luciferase reporters fused to one of two histone UTRs (**Figure 2E, Supplemental Figure S2F**). Transfection of synthetic tRF-GG drove increased luciferase activity (30%, p = 0.0002), while tRF-GG inhibition resulted in decreased luciferase levels (with values ranging from 14% to 32% in five separate experiments - each in at least triplicate - with p values ranging from 0.038 to 0.000019). tRF inhibition had no effect on a stable ES cell line carrying the wild-type luciferase reporter (data not shown), indicating that the histone 3’ UTR is necessary to confer regulation. Moreover, loss of histone reporter activity was specific to tRF-GG inhibition, as it was not observed in response to four other tRF-directed antisense LNA oligonucleotides (**Supplemental Figure S2G**). We conclude from these data that tRF-GG regulates histone mRNA abundance via the histone 3’ UTR.

### tRF-GG affects histone expression and MERVL repression via control of U7 noncoding RNA

Proper biogenesis of the histone mRNA involves a complex ribonucleoprotein assembly of 3’ UTR-associated proteins, as well as the noncoding U7 RNA which directs UTR processing via basepairing to the HDE of the histone 3’ UTR (Marzluff and Koreski, 2017). Intriguingly, in addition to downregulation of histone genes, we noted that the other consequence of tRF-GG KD in both human and mouse ES cells was decreased expression of several major classes of noncoding RNA, including snoRNAs, scaRNAs, and, to a lesser extent, various spliceosomal ncRNAs (**Figures 3A-B**, **Supplemental Figure S4**, **Supplemental Table S2**). Notably, all of these RNAs share a common biogenesis pathway with U7, as these RNAs are all processed in Cajal bodies (Gall, 2000; Machyna et al., 2013; Wu and Gall, 1993). To determine whether tRF-GG also affected levels of U7 RNA (which was not detected in our RNA-Seq data, potentially as a consequence of its secondary structure), we assayed U7 levels in tRF-GG KD and overexpression cells by Northern blotting (**Figure 3C**) and q-RT-PCR (**Supplemental Figures S4B, D**). Consistent with the effects of tRF-GG manipulation on other noncoding RNAs, we find that inhibition of tRF-GG led to reduced U7 expression, while transfecting cells with the synthetic tRF-GG oligo supported higher expression of U7. Together, these findings reveal a conserved role for tRF-GG in promoting noncoding RNA production associated with Cajal bodies, and suggest that its effects on the histone 3’ UTR result from altered U7 levels.

**Figure 3.**
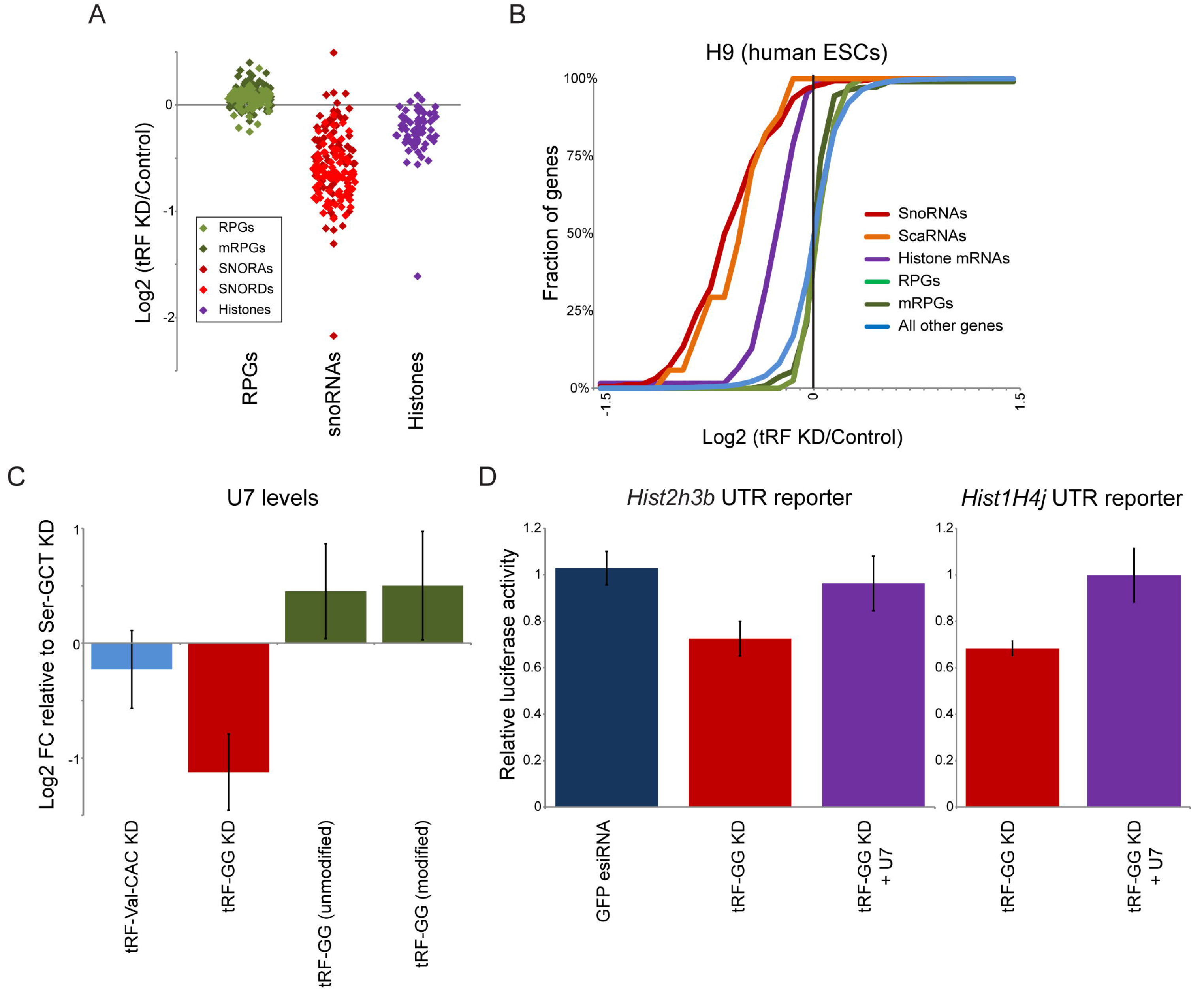
tRF-Gly-GCC supports production of U7 and other noncoding RNAs. A) Effects of tRF inhibition on various gene families in human H9 ES cells. Individual dots show individual species for the indicated families, illustrating the widespread downregulation of histone and snoRNA genes in response to tRF-Gly-GCC inhibition. Ribosomal protein genes are shown as a representative highly-expressed but tRF-insensitive gene family for comparison. Importantly, the role for tRF-GG in control of histone mRNAs and various noncoding RNAs was previously missed because prior analyses focused on polyadenylated mRNA abundance, while the current study analyzed rRNA-depleted total RNA samples (**Methods**). See also **Supplemental Figure S4**. B) CDF plots showing the distribution of tRF effects on the indicated gene families, as in **Figure 2C**. C) Manipulating tRF-GG levels affects U7 ncRNA production. ES cells were transfected either with LNA antisense oligos targeting tRF-Ser-GCT, tRF-Val-CAC, or tRF-GG, or with synthetic tRF-GG oligos either bearing appropriate modified nucleotides (modified) or lacking these modifications (unmodified). U7 levels were quantitated by Northern blot (n=4), and normalized relative to 5S rRNA levels. Change in U7 levels is expressed relative to tRF-Ser-GCT inhibition, revealing a significant (p=0.03) decrease in U7 levels in response to tRF-GG inhibition, as well as modestly increased U7 levels in tRF-GG-supplemented cells. See also **Supplemental Figure S5**. D) Effects of tRF-GG KD on histone 3’ UTR reporters are suppressed by supplementation with additional U7 snRNA. ES cells were transfected with the LNA antisense to tRF-GG, with or without additional in vitro-synthesized U7 RNA. Effects of tRF-GG KD were significant (p = 0.0039 and 0.00013 for H3 and H4 reporters, respectively), while tRF-GG KD + U7 was statistically indistinguishable from control (p=0.24 and 0.48, respectively).

The hypothesis that tRF-GG control of U7 levels is responsible for changes in histone and MERVL expression makes the prediction that manipulating U7 levels should suppress tRF-GG effects on histone expression and MERVL targets. We therefore transfected our histone 3’ UTR mESC line with the anti-tRF-GG LNA - which results in decreased U7 levels - with or without supplementation of additional U7 RNA, and assayed luciferase activity and MERVL target gene expression by qRT-PCR.

Restoring U7 levels in tRF-GG KD cells reversed the inhibition of histone expression in these cells as assayed by both luciferase reporters (**Figure 3D**) and by qRT-PCR (**Supplemental Figure S5A**). The converse also held true - antisense oligonucleotides directed against U7 were able to reverse the increase in histone levels in ES cells transfected with excess tRF-GG (**Supplemental Figure S5A**). Importantly, restoring histone mRNA levels via U7 replenishment in tRF-GG-inhibited cells was able to partially suppress the transcriptional derepression of MERVL-linked genes as assayed both by qRT-PCR and using a MERVL-driven fluorescent reporter cell line (**Supplemental Figures S5B-D**). This supports a pathway in which MERVL repression is downstream of tRF-GG-mediated histone expression, rather than being secondary to the tRF’s effects on snoRNA or other noncoding RNA production.

### hnRNPF/H are tRF-GG binding proteins and are robust repressors of the 2C-like state

Finally, we turn to the question of how tRF-GG alters noncoding RNA production. To identify direct binding partners of tRF-GG, we used a 3’-biotinylated tRF-GG to isolate candidate tRF-binding proteins from mESC whole cell extracts. tRF-GG, but not the unrelated tRF-Lys-CTT oligo, pulled down a protein of ~50 kD (**Figure 4A**). We carried out four replicate mass spectrometry analyses, in three cases analyzing bulk tRF-GG or tRF-Lys-CTT pulldowns, and in one case first gel-purifying the tRF-GG-enriched ~50 kD band. Together, these efforts identified several potential binding partners enriched in tRF-GG pulldowns relative to the control tRF-Lys-CTT pulldown (**Supplemental Table S3**). Based on follow-up functional studies of several top candidates (see below), we focus on the highly homologous pair of heterogeneous nuclear ribonucleoproteins (hnRNP) hnRNPF and hnRNPH (**Figures 4B-C**). These RNA-binding proteins function redundantly *in vivo*, and will therefore be referred to below as “hnRNPF/H”. Using an antibody that detects both hnRNPF and hnRNPH, we confirmed the interaction between tRF-GG and hnRNPF/H by Western blotting - hnRNPF/H was robustly detected in tRF-GG pulldowns, with only modest hnRNPF/H levels detected following tRF-Lys-CTT pulldown (**Figure 4D**). Furthermore, we expressed and purified hnRNPH1 protein, and carried out quantitative gel shift and fluorescence polarization analyses of hnRNPH1 binding to a fluorescently-labeled tRF-GG oligonucleotide. Both assays revealed specific binding between hnRNPH1 and tRF-GG, in contrast to the nearly undetectable binding observed for tRF-Lys-CTT (**Figure 4E-H**, **Supplemental Figure S6**). We note that the apparent K_d_ for hnRNPH1 binding to tRF-GG is roughly 5-fold weaker (K_d, app_ ~230 nM) than that for a positive control - a previously-described hnRNPF/H binding site identified in the SV40 pre-mRNA (Alkan et al., 2006) (K_d_ ~50 nM). Given that hnRNPF/H are abundant enough to regulate targets bound with 50 nM affinity, and the potential for tRF-GG to occur at much higher abundance than typical pre-mRNA targets, the apparent K_d_ of hnRNPH1 for tRF-GG is well within a plausible physiologically-functional range.

**Figure 4.**
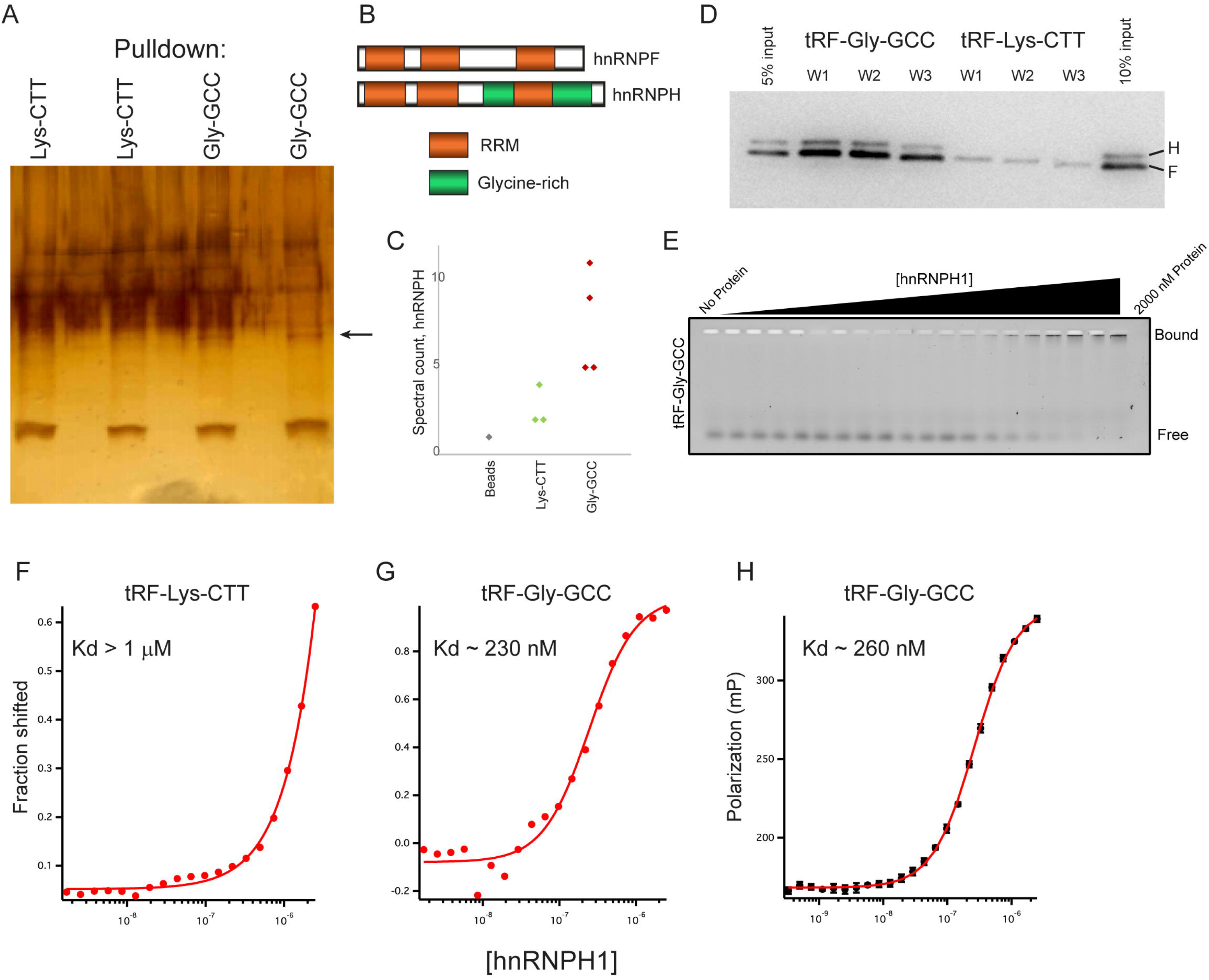
tRF-Gly-GCC binds to hnRNPF/H. A) Biotin-oligo pulldowns from murine ES cell extracts. Silver stained gel shows two replicates each for pulldowns using biotin-tRF-GG or biotin-tRF-Lys-CTT, as indicated. Arrow indicates ~50 kD band enriched in tRF-GG pulldowns. B) Domain architecture of hnRNPF and hnRNPH. C) Peptide counts for hnRNPH in control, tRF-Lys-CTT, or tRF-GG pulldowns. D) Western blots show hnRNPF/H recovery following tRF-GG or tRF-Lys-CTT pulldown. Pulldowns were washed 4 × 3 minutes with 50 mM Tris pH 8.0 supplemented with 100 mM (W1), 250 mM (W2), or 500 mM (W3) NaCl. E) Gel shift analysis of hnRNPH1 binding to tRF-GG. A synthetic oligonucleotide corresponding to tRF-Gly-GCC (GCAJULGUGGUUCAGUGGDAGAAUUCUCGC) was labeled at the 3’ end using fluorescein 5-thiosemicarbazide, then incubated at 3 nM in equilibration buffer (0.01% Igepal, 0.01 mg/ml carrier tRNA, 50 mM Tris, pH 8.0, 100 mM NaCl, 2 mM DTT) for 3 hours along with increasing concentrations of purified hnRNPH1 protein from 1.35 nM to 2000 nM. See also **Supplemental Figure S6**. F-G) Fit of gel shift binding data for tRF-Lys-CTT and tRF-Gly-GCC. Fitting the binding data yields an estimated Kd of hnRNPH1 of ~230 nM for tRF-Gly-GCC, and >1 μM for tRF-Lys-CTT. H) Fluorescence polarization data for hnRNPH1 incubations with labeled tRF-Gly-GCC. Polarization values against the protein concentrations are fit to the Hill equation using the Igor Pro software.

Do hnRNPF/H share any of the *in vivo* functions we identified for tRF-Gly-GCC? To determine potential roles for hnRNPF/H in histone gene regulation and MERVL repression, we carried out RNA-Seq in ES cells subject to double knockdown of both hnRNPF/H - we note that hnRNPF and H can compensate for one another; for example, both hnRNPH RNA and protein are upregulated in hnRNPF knockdown cells (not shown). Double knockdown of hnRNPF/H resulted in dramatic alterations in expression of several hundred genes (**Figure 5A**), including significant downregulation of developmentally-relevant genes (*Sfrp4*, *Otx2*, *Dact2*, *Spry1/4*, *Gbx2*, *Notum*, *Notch4*, *Sall1, Fgf15*, *Tdgf1*, *Inhbb*, *Ltbp3*, *Fgf4*, *Pou4f2*, *Prdm14*, *Lefty2*, *Bmp4*, etc.). The misregulation of this wide array of differentiation-associated genes presumably results from the recently-discovered role for hnRNPF/H in alternative splicing of Tcf3, a key transcription factor involved in stem cell maintenance and differentiation (Yamazaki et al., 2018).

**Figure 5.**
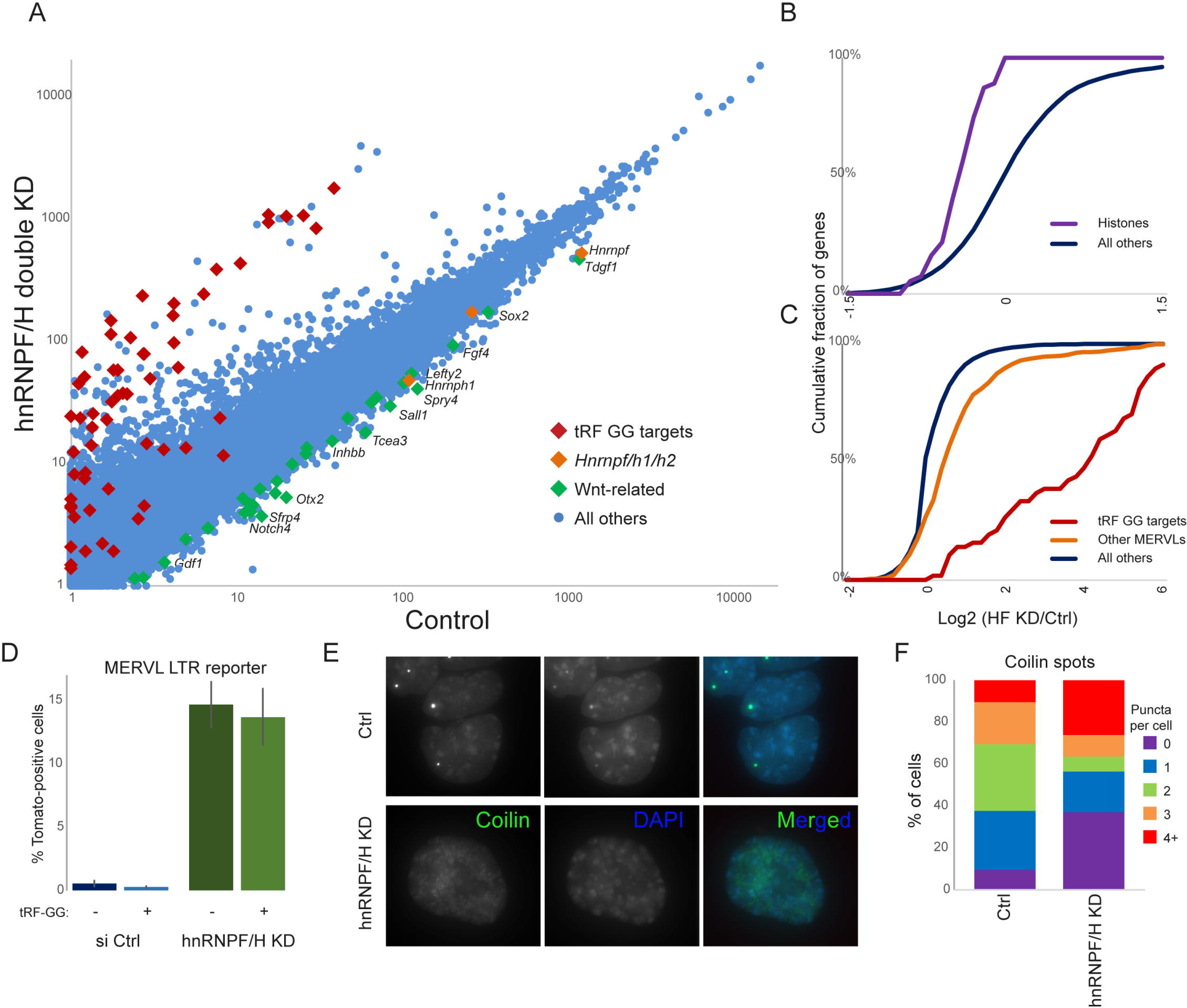
hnRNPF/H represses the MERVL program. A) Changes in the ES cell transcriptome following hnRNPF/H knockdown. Scatterplot shows mRNA abundance compared between control KD cells (x axis) and hnRNPF/H KD cells (y axis). Data are shown as median mRNA abundance for three biological replicates (**Supplemental Table S4**). B) hnRNPF/H KD results in histone mRNA downregulation. Cumulative distribution plot shows log2 fold change (hnRNPF/H KD/Ctrl) for histone genes, and all other genes, as indicated. C) Effects of hnRNPF/H KD on expression of the MERVL program. Cumulative distribution plots show log2 fold change (hnRNPF/H KD/Ctrl) for tRF-GG target genes (Sharma et al., 2016)), other genes previously associated with the MERVL program (Macfarlan et al., 2012), and all other genes, as indicated. D) hnRNPF/H suppresses ES cell entry into the 2C-like state. ES lines carrying a MERVL LTR-driven tdTomato were subject to control or hnRNPF/H KD, with bars showing mean +/- standard deviation (n=5 replicates) of the percentage of Tomato-positive cells. E) hnRNPF/H is required for normal Cajal body morphology and gross chromatin architecture. Panels show typical images for the Cajal body marker coilin (green) and DAPI (blue), in control or hnRNPF/H KD ES cells. See also **Supplemental Figure S7A**. F) Quantitation of Cajal body number per cell. Stacked bars show the percentage of cells exhibiting 0 through 4+ coilin-positive puncta, as indicated. Data represent the average of two replicate transfections, with 85 total control and 98 hnRNPF/H KD cells quantitated across the two experiments.

Consistent with the possibility that tRF-GG functions through hnRNPF/H binding, we find that hnRNPF/H also affected the same groups of genes regulated by tRF-GG, with hnRNPF/H knockdown resulting in downregulation of histone genes and a dramatic derepression of the MERVL program (**Figures 5A-C**). Using a MERVL reporter ES cell line (Macfarlan et al., 2012), we find that hnRNPF/H KD led to a ~30-fold increase in Tomato-positive cells (**Figure 5D, Supplemental Figure S7A**), confirming the derepression of the MERVL program observed in the RNA-Seq dataset. Importantly, transfection of synthetic tRF-GG had no effect on MERVL repression in the absence of hnRNPF/H (**Figure 5D**), consistent with the hypothesis that tRF-GG acts by binding these proteins. Next, given the general role for tRF-GG in supporting the output of multiple Cajal body products, we examined Cajal body morphology in hnRNPF/H KD cells, using the well-known Cajal body marker coilin (Gall, 2000). In contrast to control ES cells, which exhibit one or two bright Cajal bodies per nucleus, we find that knockdown of hnRNPF/H leads to more diffuse coilin staining (**Figures 5E-F, Supplemental Figure S7A**). Moreover, DAPI staining was clearly distinctive in hnRNPF/H KD cells, with the typical discrete chromocenters being replaced by more diffuse “lumpy” DAPI staining (**Figure 5E**), potentially secondary to the dramatic changes in histone expression in these cells. Taken together, our data reveal a novel role for hnRNPF/H in supporting normal Cajal body function in mouse embryonic stem cells, with downstream consequences for histone expression and chromatin-mediated repression of MERVL elements.

## DISCUSSION

Here, we investigated the mechanism by which tRF-Gly-GCC represses MERVL-associated gene expression in ES cells and preimplantation embryos. Surprisingly, we find that tRF-GG indirectly represses MERVL-driven transcription as a downstream consequence of its effects on noncoding RNA biogenesis. tRF-GG regulates a cascade of events through its role in supporting U7 snRNA levels, which in turn enhances expression of canonical histone genes, ultimately resulting in increased heterochromatin-dependent repression of LTR-associated genes (**Figure 6**).

**Figure 6.**
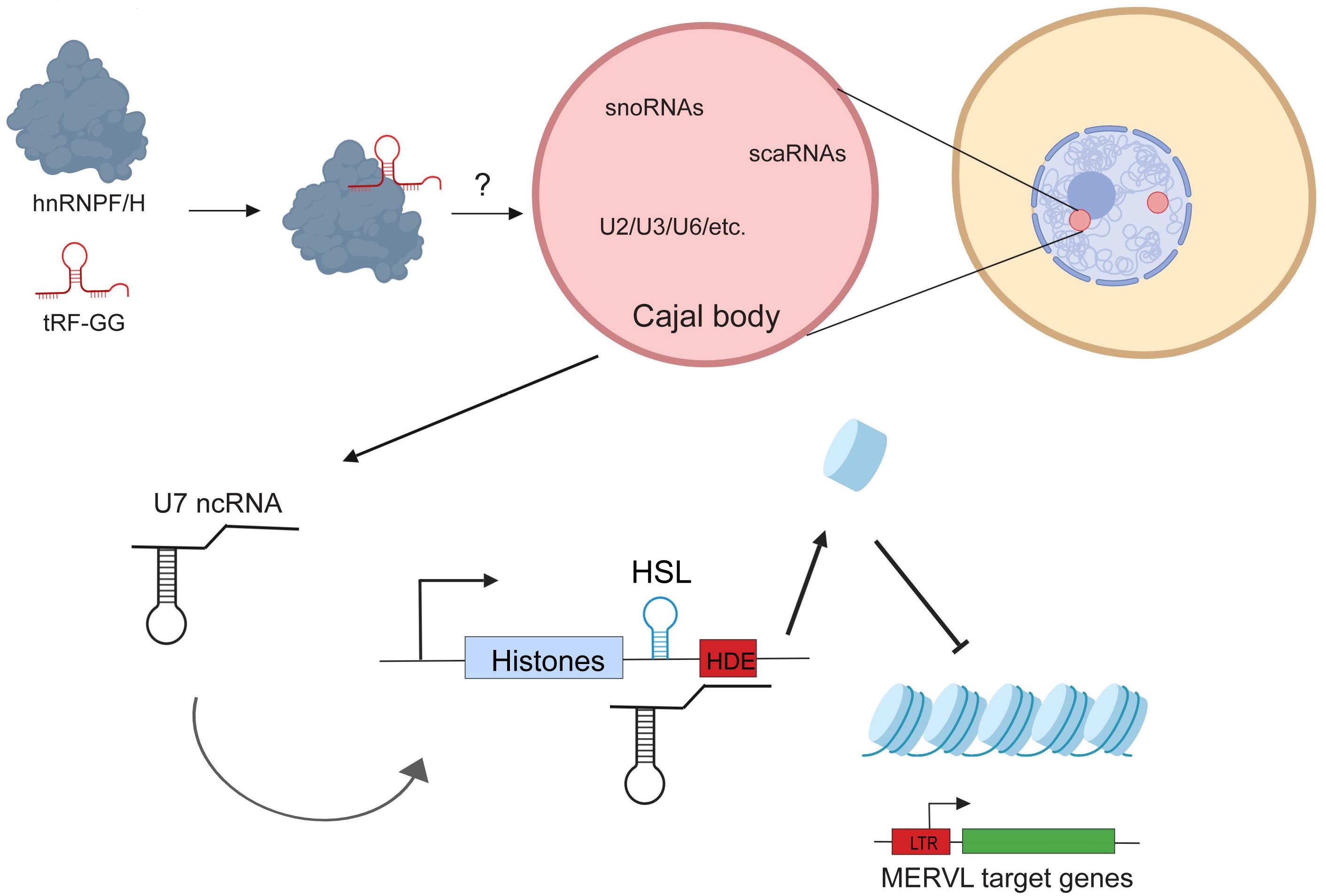
Schematic of proposed mechanism for tRF-GG function. Together, our data support a model in which 5’ tRF-Gly-GCC binds to hnRNPF/H and serves to support production of a variety of noncoding RNAs in Cajal bodies. Central to the current study is control of U7 ncRNA production, which plays a central role in processing of the histone 3’ UTR via base-pairing to the histone downstream element (HDE). Altered expression of histones then leads to downstream effects on the expression of MERVL-associated genes in murine embryonic stem cells and preimplantation embryos. The precise mechanism by which tRF-GG supports hnRNPF/H function remains to be elucidated - tRF-GG could stabilize hnRNPF/H leading to increased functional protein levels, or hnRNPF/H and tRF-GG could function together in a complex as depicted here.

Our data demonstrate the effects of tRF-GG on MERVL target genes are mediated by altered histone gene expression. We document effects of tRF inhibition and tRF “overexpression” on histone expression by q-RT-PCR, by deep sequencing, and by quantitative Western blot, we confirm the expected downstream effects on chromatin compaction by ATAC-Seq, and finally we show that tRF regulation of histone genes can be recapitulated using two distinct histone 3’ UTR reporters. Moreover, the effects of tRF-GG manipulation - both inhibition and overexpression - on histone regulation and on MERVL target repression can be suppressed by appropriate manipulation of the U7 noncoding RNA. Our data argue that, rather than being directly targeted by tRF-GG via sequence homology between tRNAs and the PBS of LTR elements, MERVL instead represents a highly sensitive reporter of global chromatin status in mouse embryonic stem cells. Further supporting this idea is the fact that histone downregulation, rather than ERV derepression, is a conserved consequence of tRF-GG inhibition in both mouse and human ES cells.

### tRF-Gly-GCC supports Cajal body output

Upstream of the histone 3’ UTR, we document a surprising and conserved role for tRF-GG as a positive regulator of noncoding RNA production, with effects of tRF manipulation on the histones resulting from altered levels of the U7 noncoding RNA. Indeed, perhaps the most unanticipated discovery described here is the finding that the 5’ tRF-Gly-GCC supports production of noncoding RNAs that are normally synthesized or processed in the Cajal body. This is demonstrated using both gain and loss of function approaches in both mouse and human ES cells. Given the wide range of functions ascribed to Cajal body ncRNAs, our data also predict that tRF-GG may affect additional downstream pathways beyond the MERVL program previously described. Confirming this prediction, we find subtle effects of tRF-GG inhibition on rRNA 2’-O-methylation (**Supplemental Figure S7D-E**), which is mediated by snoRNAs, as well as a small global increase (~5%) in intron retention in tRF-inhibited cells (**Supplemental Table S1**), consistent with the observed decrease in spliceosomal RNAs. Thus, fine-tuning of Cajal body output by tRF-GG affects cellular functions ranging from splicing to translation to global chromatin packaging.

### hnRNPF/H are novel regulators of the 2C-like state

What is the proximate mechanism by which tRF-GG supports production of Cajal body RNAs? We identify a strong candidate for a relevant effector protein, finding that the closely related hnRNPF and H proteins bind directly to tRF-GG in extracts and *in vitro*. Moreover, functional studies reveal that hnRNPF/H and tRF-GG exhibit heavily overlapping regulatory roles in vivo, identifying a novel and surprising role for hnRNPF/H in control of the MERVL program in mES cells. Indeed, hnRNPF/H represent the strongest repressors of the 2C-like state yet observed, as the ~30-fold derepression of the 2C-like state in response to hnRNPF/H KD is comparable to, and in fact more dramatic than, that previously observed following Chaf-1 or Ubc9 knockdown (Cossec et al., 2018; Ishiuchi et al., 2015).

hnRNPF/H are RNA-binding proteins with relatively well-characterized roles in mRNA splicing (Wang et al., 2012; Wang et al., 2007; Yamazaki et al., 2018), and our RNA-Seq analysis of hnRNPF/H KD cells reveal hundreds of altered splicing events (see **Supplemental Figures S7B-C** for examples), consistent with previous studies (Yamazaki et al., 2018). How do these changes in RNA splicing - or, alternatively, some unrelated activity of hnRNPF/H - ultimately drive repression of the MERVL program? Given the dramatic changes in Cajal body morphology documented in hnRNPF/H KD cells (**Figure 5**), we favor the hypothesis that one or more hnRNPF/H-regulated transcripts plays a key role in Cajal body function, with altered U7 RNA production altering histone production and thereby driving changes in the highly heterochromatin-sensitive MERVL program. A number of specific target(s) of hnRNPF/H could be responsible for supporting normal Cajal body biogenesis, as a large number of chromatin and RNA-binding proteins (such as Hnrnpa2b1) with potential roles in Cajal bodies exhibit altered splicing patterns in hnRNPF/H-depleted cells. Given that tRF-GG only affects a small subset of hnRNPF/H functions - tRF-GG inhibition has no effect on *Tcf3* splicing (**Supplemental Figure S7B**), for example - we speculate that the hnRNPF/H targets responsible for Cajal body biogenesis are the most sensitive to subtle changes in hnRNPF/H function. It will be interesting in future studies to determine the mechanism by which tRF-GG enhances the function of hnRNPF/H - whether tRF-GG stabilizes hnRNPF/H for example, or whether hnRNPF/H functions in complex with tRF-GG at a subset of targets.

### Implications for tRF-Gly-GCC function in the early embryo

We finally turn to the question of the physiological contexts in which tRF-Gly-GCC is likely to play an important role in cellular function. In typical somatic tissues, tRNAs are cleaved in response to a variety of stress conditions; for example tRNA cleavage occurs in response to arsenite treatment of neurons to produce tRNA fragments which help to direct global translational downregulation (Ivanov et al., 2011). Curiously, beyond the case of stress-dependent tRNA fragment production, it is increasingly clear that tRNA cleavage occurs commonly in the germline of multiple organisms even under apparently stress-free growth conditions (Couvillion et al., 2010; Peng et al., 2012), and tRF-GG is one of the most abundant small RNAs present in mammalian sperm and delivered to the zygote upon fertilization. Although we previously showed that manipulating tRF-Gly-GCC levels in the zygote altered expression of ~50 MERVL-associated transcripts, these results were based on single-embryo mRNA-Seq which only reports on transcripts bearing a polyA tail. However, the wide variety of noncoding RNAs affected by tRF-Gly-GCC are predicted to affect an extensive list of additional cellular processes from ribosome biogenesis (snoRNAs) to splicing (scaRNAs and U RNAs) to global chromatin compaction (U7). How modulation of these various functions by tRF-GG delivery to the zygote affects later processes during preimplantation development, and whether any of these alterations have lasting consequences for offspring phenotypes, will be of great interest.

## ACKNOWLEDGEMENTS

We thank Zbig Dominski for the U7 expression vector. We thank Sean Ryder and members of the Rando lab for critical reading of the manuscript and insightful discussions. This work was supported by NIH grant R01HD080224. A.B. was supported by a fellowship from the Human Frontier Science Programme (LT000857/2015-L).

## METHODS

### Mouse ES cell culture and transfections

All murine ES cell lines were grown in DMEM (Gibco) containing 10% fetal bovine serum and leukemia inhibitory factor (Serum+LIF culture conditions) and all transfections were carried out using OptiMEM and Lipofectamine 2000 (Invitrogen) according to the manufacturer’s instructions at splitting, unless otherwise specified. Inhibition of tRF-Gly-GCC function in mESCs was performed as described in Sharma et al, 2016. Controls included Lipofectamine 2000 only (Mock), anti-GFP esiRNA transfections and/or a scrambled anti-tRF-GG LNA oligo. Transfection of various 3’tRFs (**Supplementary Table S5**) was performed as for the 5’tRF-GG. Concentrations tested were 5 ng and 100 ng.

U7 rescue experiments were performed by supplementing the tRF-Gly-GCC inhibition transfection reaction with 50 ng of in vitro synthesized U7 snRNA per 3×10^5 cells. Mouse U7 small nuclear RNA was generated through in vitro transcription from pGEM-Teasy-U7 plasmid (generous gift from Z.Dominski) using mMessage mMachine T7 kit (Ambion) following plasmid linearization with HindIII restriction enzyme. Human U7 was cloned from hESC cDNA into the pGEM-Teasy plasmid and used as in mouse experiments. Media was changed after 16 hrs and cells were allowed to grow for additional 32 hrs. Cells were processed for various experiments at the end of the 48-hour period. U7 knockdown was achieved by transfecting 10 ng of modified anti-sense oligonucleotides targeting U7 (**Supplementary Table S5**) synthesized with phosphorothioate linkages.

Double hnRNPF/H knockdown was performed by transient transfection of 20 pmols of siRNAs against mouse hnRNPF and hnRNPH transcripts, respectively (sc-38273 and sc-35580, Santa Cruz Biotechnologies) in 12-well format. 40 pmols of siRNA-A (sc-37007, Santa Cruz Biotechnology) was used as knockdown control. Efficiency of knockdown was validated by Western Blotting and RNA-sequencing. Cells were collected 48 hours after transfection for various downstream experiments.

### Cell culture (H9)

Line H9 (WA09) human embryonic stem cells were cultured on Matrigel (Corning) in mTESR1 media (Stem Cell Technologies) in 5% CO2 at 37°C. Nucleofection of oligos was done using Human Stem Cell Nucleofector Kit 1 (Lonza Bioscience), according to manufacturer’s instructions. 2 ng of LNA was nucleofected by 24-well plate. Nucleofection efficiency was checked using pEGFP plasmid control, with successful experiments having more than 80% GFP+ cells. Cells were harvested 12hours post-nucleofection for RNAseq and qRT-PCR.

### Metabolic labeling

E14s were labeled in 500mM 4sU containing media for 15 or 30mins, then RNA was isolated using Trizol and isopropanol precipitation. 50 μg of total RNA was mixed with 0.2mg/ml of EZ-Link Biotin-HPDP (Themo Fisher) in 500 μL reaction then incubated for 2hr at 37°C in a shaking thermomixer (750 rpm). Biotinylated RNA was then extracted using phenol:chloroform:isoamyl alcohol (PCI) with phaselock gels and precipitated using isopropanol. RNA pellet was resuspended in 10μL water, and mixed with 30μL of washed Dynabeads MyOne Streptavidin C1 beads (Invitrogen) in binding buffer (10 mM Tris-HCl pH7.5, 300 mM NaCl, 0.1% Triton-X). The slurry was rotated for 20 mins at room temperature to immobilize biotin-tagged RNA, then placed on magnetic stand and washed with 500μL high salt buffer (50 mM Tris-HCl, 2M NaCl, 0.5% Triton-X). The supernatant from the first high-salt wash is the unlabeled total RNA. Beads were then stringently washed in high salt buffer, then two times in binding buffer, then once in low salt buffer (5 mM Tris-HCl, 0.1% Triton-X). The biotin-tagged RNA were extracted from the beads with 100 mM DTT at 65°C for 5 mins twice. Finally, labeled and unlabeled fractions were PCI extracted and RNA was isopropanol precipitated, and used to construct RNA-seq libraries.

### RNA-seq

5 μg of total RNA was depleted of ribosomal RNA using Ribo-Zero rRNA Removal Kit (Human, Mouse, Rat, Illumina). Less total RNA was used as input from metabolic labeling experiments for both 4sU labeled and unlabeled fractions. Illumina deep sequencing compatible libraries were constructed from rRNA-depleted RNA using an optimized version of a protocol described by Heyer et al. (Heyer et al., 2015), adding a purification using the RNA Clean and Concentrator (Zymo Research) in between procedures. Ribosome profiling data was published previously (GSE74537), with libraries constructed using the same procedure. Libraries were quantified, multiplexed and either single-end or paired-end sequenced on Illumina NextSeq 500 sequencer.

### ATAC-Seq

E14 mES cells were transfected with antisense tRF-Gly-GCC LNA or mock-transfected and grown for 24 or 48 hours prior to harvesting and counting. ATAC-seq protocol was done essentially the same as described in Buenrostro et al. Briefly, after titration, 4 μL of TDE1 was determined as sufficient for 50000 cells. Tagmented DNA was amplified using Kapa HiFi Hotstart polymerase, and libraries were cleaned up using Ampure XT DNA beads. Libraries were quantified, multiplexed and paired-end sequenced on an Illumina NextSeq 500 sequencer.

### Ribometh-seq

Ribometh-seq was done essentially as previous described (Marchand et al., 2016). Briefly, 500 ng of total RNA was fragmented with 100mM alkaline solution at 95 °C for 10 mins. Fragmented RNA was then 3’ dephosphorylated with T4 PNK for one hour, followed by 5’ phosphorylation using T4 PNK in the presence of ATP for one hour. RNA was concentrated using Zymo RNA Clean and Concentrator, and libraries were constructed from 100 ng of prepared RNAs using NEBNext Small RNA Library Prep Kit following manufacturer’s instructions. Ribometh-seq libraries were single-end sequenced on an Illumina NextSeq 500 sequencer.

### Deep sequencing data analysis

RNA-seq libraries were demultiplexed using Novobarcode (v3.02.08). Single end libraries were trimmed of 3’ adapters using Fastx-toolkit (v0.0.14). Quantification was done using RSEM (v1.2.29) to RefSeq GTF annotation, mapped with Bowtie (v1.0.0) to mm10 using default parameters.

For splicing analysis, reads were mapped using STAR aligner (v2.5.3a) using the “two-pass” method. Briefly reads were mapped to mm10 genome using default settings, then reads were remapped to the collective set of junctions predicted from the first pass. Alternative splicing was analyzed using JUM (2.0.2, (Wang and Rio, 2018)). Alternative splicing events with a p-value less than 0.05 were extracted for further analysis.

ATAC-seq libraries were mapped to mm10 using Bowtie2 (v2.3.2) with the following parameters: −D 15 −R 2 −N 1 −L 20 −i S,1,0.50 --maxins 2000 --no-discordant -- no-mixed. Fragment lengths were separated using Python, and coverage of reads in various chromatin states was analyzed in R (v3.4.1) using data from ChromHMM (https://github.com/guifengwei/ChromHMM_mESC_mm10). All coverage data was normalized by global read depth prior to further analysis. Circos plots (v0.69-2) were generated from coverage data calculated by Bedtools (v2.25.0).

Ribometh-seq libraries were mapped using Bowtie2 and the 5’ locations of the forward read was quantified using Bedtools.

### qRT-PCR and semi-quantitative PCR

For mESCs, RNA was isolated using Trizol (Ambion) according to the standard protocol and the samples were treated with TurboDNase to eliminate genomic DNA contamination. Following TurboDNase treatment, RNA was purified using Zymo RNA clean and concentrator 5 kit, quantified on Nanodrop and RT reaction was performed using SuperScript IV RT kit (Invitrogen) according to the manufacturer’s protocol. Obtained cDNA was diluted 2X for MERV-L target gene qPCR) and 10X for histone and β-actin amplification (based on the standard curves obtained for the primers used) and Kapa SYBR Fast Universal Mastermix was used in all q-RT-PCR reactions. The amplification conditions were: 95C for 3 minutes followed by 40 cycles of 95C for 5s and 60C for 20 s (with a plate-reading step between each cycle) based on the previously reported conditions for histone qPCR. All the reactions were read on the BioRad CFX96 qPCR detection machine. The same protocol, with human-specific primers, was performed for H9 human cells.

For semiquantitative PCRs to confirm candidate alternative splicing events, primers were designed in flanking exons. The number of PCR cycles were empirically determined to avoid saturation of PCR amplicons.

### Western Blotting

Mock or tRF-GG inhibited cells were grown as before and 48 hours after transfection, cells were trypsinized and counted. A defined number of cells (usually 50000) was spun from each group, washed in PBS, pelleted and lysed directly by boiling at 100 °C for 15 minutes in 2X Laemmli Sample buffer containing β-mercaptoethanol. 2X serial dilutions of the cell lysates were loaded onto a 15% SDS-polyacrylamide gel. Following protein resolution on the gel, proteins were transferred to a nitrocellulose membrane for 1 hour at 100 V through wet transfer. Membrane was blocked in 5% milk in TBSt for at least an hour prior to the addition of primary antibodies overnight at 4 °C. Primary antibodies used were anti-histone H3 (Millipore, #07-690), anti-histone H4 (Millipore, #05-858), anti β-actin (abcam ab8224) and anti-GAPDH (abcam ab9485). Membranes were then washed 3X in TBSt for 15 minutes at room temperature. Membrane was incubated with secondary antibody in 5% milk for 1 hour at room temperature. Secondary antibodies used were anti-mouse IgG, HRP-linked (Cell Signaling, #7076S) and anti-rabbit IgG, HRP-linked (Cell Signaling 7074S). Following 3 additional TBSt washes, the substrate for HRP was added (Amersham ECL Western Blotting Analysis System, GE) and the blots were exposed on the Amersham Imager 600 machine.

### Northern Blotting

DNA probes for snoRNAs were ordered from IDT (**Supplementary Table S5**) and 5’ end-labeled with [γ-^32^P] using PNK (NEB). U7 was present in very low abundance and had to be probed using a full-length [γ-^32^P]C labeled RNA probe synthesized from a pGEM plasmid containing a cloned mouse U7. 2ug of total RNA was separated on a 6% urea PAGE gel. RNA was transferred onto a Hybond-N+ membrane (GE Healthcare) by semi-dry transfer method (BioRad) in SSC buffer. RNA was then crosslinked to the membrane at 254nm. Probes were then hybridized to RNA overnight with agitation at 68C. Membranes were then washed, and exposed to BioMax film (Kodak) for up to three days. Data was quantified using ImageJ.

### Embryo microinjection studies

Zygotes were generated through IVF, as described previously (Sharma et al., 2016). After IVF, embryos were washed and placed in KSOM for culturing for 2 hours to recover. Following this, zygotes were washed 2X in M2 media and microinjected with a) control (H3.3-GFP mRNA at 100 ng/μl) and b) tRF-modified (H3.3-GFP mRNA+ mod-tRF-GG at 200 ng/μl). After micromanipulation, zygotes were placed back in KSOM for overnight culture and the following day (24h post-IVF) fluorescence was checked in 2-cell stage embryos. GFP-positive embryos were washed 2X in M2 media and their zona pellucida was removed by acid tyrode solution. Embryos were then neutralized by 2 washes in M2, and mouth pipetted into 5 μl ATAC-lysis buffer (containing 1% TritonX, without NP-40) on ice. 15 μl of tagmentation reaction mix was added (11.25 μl TD buffer, 1.5 μl TDE1, 2.25 μl H20) and the reaction was incubated for 30 minutes at 37C in a thermoblock with intermittent shaking. Tagmentation reaction was stopped by the addition of EDTA (25 mM final) and incubation at 50 °C for 10 minutes. Prior to PCR amplification, reaction was supplemented with MgCl2 and the rest of the protocol was performed as described for ATAC-seq in cultured cells. After amplification, libraries were quantified by Qubit DNA High Sensitivity Assay, concentration was adjusted between control and experimental groups, and qPCR was performed using Kapa SYBR-FAST Universal qPCR mix on BioRad CFX96 Real-Time System. Amplification conditions were: 95 °C 3 minutes (initial denaturation), followed by 40 cycles: 95 °C for 10 seconds, 60 °C for 15 seconds, 72 °C for 15 seconds. The efficiency of primers for MERV-L and tubulin was validated using previously sequenced ATAC-Seq libraries generated from mouse embryonic stem cells.

### Histone 3’ UTR luciferase reporter assay

Histone H3b (*Hist2h3b*) and histone H4j (*Hist1h4j*) 3’UTR sequences (~300 nucleotides) were cloned into the PsiCheck 2 vector downstream of the Renilla luciferase coding sequence, between XhoI and NotI restriction sites. As PsiCheck2 vector does not encode for a eukaryotic selectable marker, cells were co-transfected with PsiCheck2-empty or PsiCheck2-Histone3’UTR together with a carrier plasmid pCDNA3.1+ Hygro and stable cell lines were generated as described in (Connelly et al., 2012). Following 7 days of Hygromycin selection, individual clones were picked, expanded and tested for luciferase expression. Based on their expression level, one clone from all cell lines was selected for subsequent experiments. Reporter ES cell lines were transfected as previously. 48 hours after transfection, cells were washed 2X with PBS and cell lysate for the luminescence reading was prepared as directed by the Dual-Luciferase Assay System (Promega). Firefly and Renilla luminescence was measured using GLOMAX 96 Microplate Luminometer, and Renilla luminescence was normalized to the internal control of Firefly luminescence.

### Cell cycle analysis

E14 mESCs were transfected with antisense 5’tRF-Gly-GCC LNA-containing oligo or mock transfected as described above. Media was changed 16 hours after transfection. After 24 hours, cells were synchronized in G1/S-phase of the cell cycle by single thymidine block (5 mM thymidine in culture media) for 16 hours. Following treatment, cells were washed 2X with PBS, and fresh culture media without thymidine was added and were allowed to progress through the cell cycle. Cells were collected by trypsinization and 2X PBS washes at time zero and every 2 hours for an 8 hour period after removal of the thymidine block. Cold 70% Ethanol was added to cells dropwise with light vortexing and cells were fixed for 30 minutes at 4 °C. Following fixation, cells were pelleted, washed 2X with PBS and treated with RNase A (final concentration 0.2 mg/ml). Cells were stained with Propidium Iodide solution (final concentration 10 mg/ml) and DNA content was analyzed using FACScan Flow Cytometer (Becton Dickinson).

### Oligonucleotide pulldowns and mass spectrometry

Mouse ES cells were washed 2X with PBS and collected by trypsinization. Around 20 million cells were lysed in NP-40 buffer (50 mM Tris pH 7.5, 0.1% NP-40, 150 mM NaCl, 1 mM CaCl2, 1 mM DTT, 40 U SuperaseIn, 1X Protease inhibitor cocktail) on ice for 15 minutes. Lysates were cleared by centrifugation for 15 minutes at 4°C and maximum speed and the supernatant was transferred to a new tube. The lysates were then divided in 2 and incubated with 500 pmols of tRF-Gly-GCC-biotin or tRF-Lys-CTT-biotin for 1 hour at room temperature with end-over-end rotation. Following the incubation, 100 μl of C1 Dynabeads were added to the reactions and incubated with end-over-end rotation for an additional hour. Captured RNPs were then washed three times each with low salt buffer (30 mM Tris-HCl pH7.5, 120 mM KCl, 3.5 mM MgCl2, 0.5 mM DTT), medium salt buffer (30 mM Tris-HCl pH7.5, 300 mM KCl, 3.5 mM MgCl2, 0.5 mM DTT) and high salt buffer (30 mM Tris-HCl pH 7.5, 0.5 M KCl, 3.5 mM MgCl2, 0.5 mM DTT) for 5 minutes at room temperature. Following the last wash, elution was performed by disrupting the biotin-streptavidin bond using 95% formamide and 10 mM EDTA at 65°C for 5 minutes. Eluate was boiled in 2X Laemmli sample buffer and loaded into 4-20% gradient polyacrylamide gel. Gels were stained by Coomassie Brilliant Blue solution for 1 hour at room temperature, then destained overnight in the destaining solution (40% MeOH, 10% acetic acid). Bands were cut from the gel and submitted for Mass Spectrometry at the UMass Medical School Mass Spectrometry facility.

### Streptavidin RNA pulldown assay

For each assay, 2.5 uM of biotin labelled RNA was incubated with streptavidin beads (Invitrogen) according to the manufacturer’s instructions. Beads were then incubated for 2 h with cellular lysate in binding buffer (0.01 mg/mL tRNA, 0.01% NP40, 0.1 mg/mL BSA, 50 mM Tris-Cl (pH 8.0), 100 mM NaCl, and 1/20 SuperaseIn (Ambion). After 2 hours of rotation at room temperature, the beads were washed with 200 μl of wash buffer (100 mM NaCl, 50 mM Tris-Cl (pH 8.0), 0.01% NP40, and 0.01 mg/mL tRNA) for 4 times. Proteins were eluted from beads with sample buffer for 5 min at 95 °C and equal amounts are run on an SDS polyacrylamide gel and analyzed by Western analysis.

### hnRNPH1 purification

The sequence encoding amino acids 1-449 of mouse hnRNPH1 was cloned into pMal-ac (New England Biolabs) downstream an N-terminal maltose-binding protein (MBP) tag and the cloned construct was transformed into BL21(DE3) cells. The cells were induced with 1 mM isopropyl 1-thio-β-D-galactopyranoside for 3 h, at 37 °C. to express the protein with an N-terminal MBP tag. The cells were lysed in 200 mM NaCl, 50 mM Tris, pH 8.8, 2 mM DTT, and EDTA-free protease inhibitor tablet. Amylose (New England Biolabs) affinity column was used for the first step of purification of hnRNPH1. Protein fractions were eluted in lysis buffer supplemented with 10 mM maltose. Fractions containing the protein were pooled and dialyzed into an S-column buffer (20 mM NaCl, 50 mM MOPS pH 6.0, 2 mM DTT). Purification was followed by HiTrap S at 4 °C. Elution of the protein fractions was achieved by a salt gradient ranging from a low salt buffer (20 mM NaCl, 50 mM Tris MOPS pH 6.0, 2 mM DTT) to a high salt buffer (1 M NaCl, 50 mM MOPS pH 6.0, 2 mM DTT). Pure fractions were dialyzed in a Q-column buffer (20 mM NaCl, 50 mM Tris pH 8.8, 2 mM DTT). Final purification was done using a HiTrap Q ion exchange column at 4 °C. Protein fractions were eluted by a salt gradient ranging from a low salt buffer (20 mM NaCl, 50 mM Tris, pH 8.8, 2 mM DTT) to a high salt buffer (1 M NaCl, 50 mM Tris, pH 8.8, 2 mM DTT). Pure fractions were determined by Coomassie-stained SDS-PAGE, and purified hnRNPH1 was dialyzed into storage buffer (25 mM Tris, pH 8.0, 25 mM NaCl, 2 mM DTT) and stored at 4 °C. Pure fractions were concentrated using an Amicon spin concentrator.

### Preparation of Fluorescently Labeled RNA

RNA oligonucleotides were 3′-end labeled with fluorescein 5-thiosemicarbazide as previously described (Pagano et al., 2011). Briefly, RNA is first oxidized with sodium periodate and then reacted it with fluorescein 5-thiosemicarbazide to form a covalent bond. Labeled RNA is then purified over a Sephadex G25 column.

### Electrophoretic Mobility Shift Assay

Electrophoretic mobility shift experiments and data analysis were carried out as previously described with a few modifications (Pagano et al., 2011). Briefly, 3 nM of labeled RNA was incubated with a gradient of hnRNPH1 concentration in equilibration buffer (0.01% Igepal, 0.01 mg/ml tRNA, 50 mM Tris, pH 8.0, 100 mM NaCl, 2 mM DTT) for 3 h. After equilibration, polarization readings were taken in a Victor plate reader. The samples were then mixed with bromocrescol green loading dye and loaded on a 5% native, slab polyacrylamide gel in 1× TBE buffer. The gels were run in 1× TBE buffer for 120 min at 120 volts and at 4 °C and then scanned using a fluor imager (Fujifilm FLA-5000) with a blue laser at 473 nm. The fraction of bound protein against the protein was fit to the Hill equation using Igor Pro software.

### Immunofluorescence

E14 mouse ES cells were grown and transfected as described above, and plated onto gelatinized coverslips. For immunofluorescence, cells were washed 2X in PBS and fixed in 4% paraformaldehyde for 20 minutes at room temperature with mild agitation. Fixed cells were permeabilized by 0.5% Triton-X solution for 20 minutes at room temperature followed by 3 washes in PBS containing 0.05% Tween 20 (PBS-Tween). Blocking was performed using 5% milk in TBSt for 1 hour at 37 °C. Cells were incubated in primary antibody in 3% BSA for 1 hour at 37 °C, followed by 3 washed with PBS-Tween. Primary antibodies used were anti-coilin (Abcam, ab210785, 1:50 dilution) and anti-hnRNPF/H (Santa Cruz Biotechnology, sc-32310, 1:500 dilution). Following the washes, cells were incubated in secondary antibody conjugated with fluorophores for 45 minutes to 1 hour at room temperature in the dark. Secondary antibodies used were Alexa Fluor 488 goat anti-mouse (1:1000 dilution), Alexa Flour 488 goat anti-rabbit (1:500 dilution). Following 3 additional washes with PBS-Tween, cells were mounted in Vectashield mounting medium containing DAPI for DNA visualization. Microscopy was performed on AxioObserver.Z1/7 microscope using 63X/1.4 NA oil objective. Images were analyzed using ImageJ software.

## SUPPLEMENTAL MATERIAL

**Table S1. Metabolic labeling of newly-synthesized RNAs**.

Deep sequencing data (normalized to ppm) for total RNA (“total”) or 4-thiouridine-labeled RNA (“4sU”) isolated from mES cells either transfected with siRNAs targeting GFP, or with an antisense LNA oligo targeting tRF-Gly-GCC. One replicate was performed with a 15 minute labeling time for 4sU, and two replicates were performed following 30 minutes of labeling.

**Table S2. Effects of tRF-Gly-GCC inhibition in human H9 ESCs**.

RNA-Seq dataset (normalized to ppm) for H9 ESCs transfected with LNA targeting tRF-Gly-GCC (n=3), or controls (n=1 using siRNAs targeting GFP, and n=2 using a scrambled anti-tRF-GG oligo control). “Genes” sheet shows data for reads mapping to Refseq, “RMSK” shows reads mapping to repeatmasker.

**Table S3. Mass spectrometry results for tRF-Gly-GCC-binding proteins**.

Data for four replicate mass spectrometry analyses of biotin-labeled oligonucleotide pulldowns from mESC extracts. Rep1 was run on the gel-purified ~50 kD band shown in Figure 4A, while the other three experiments were carried out on unfractionated bulk pulldown material using either beads alone, tRF-Lys-CTT, or tRF-Gly-GCC.

**Table S4. Effects of hnRNPF/H double knockdown on gene expression in mESCs**.

RNA-Seq data (normalized to ppm) for control and hnRNPF/H knockdown ESCs. Knockdowns were performed for hnRNPH alone, or both hnRNPF and H, as indicated.

**Table S5. Oligonucleotides used in this study**.

Oligonucleotide sequences for synthetic tRFs and for RT-PCR primer pairs used in this study.

**Figure S1.**
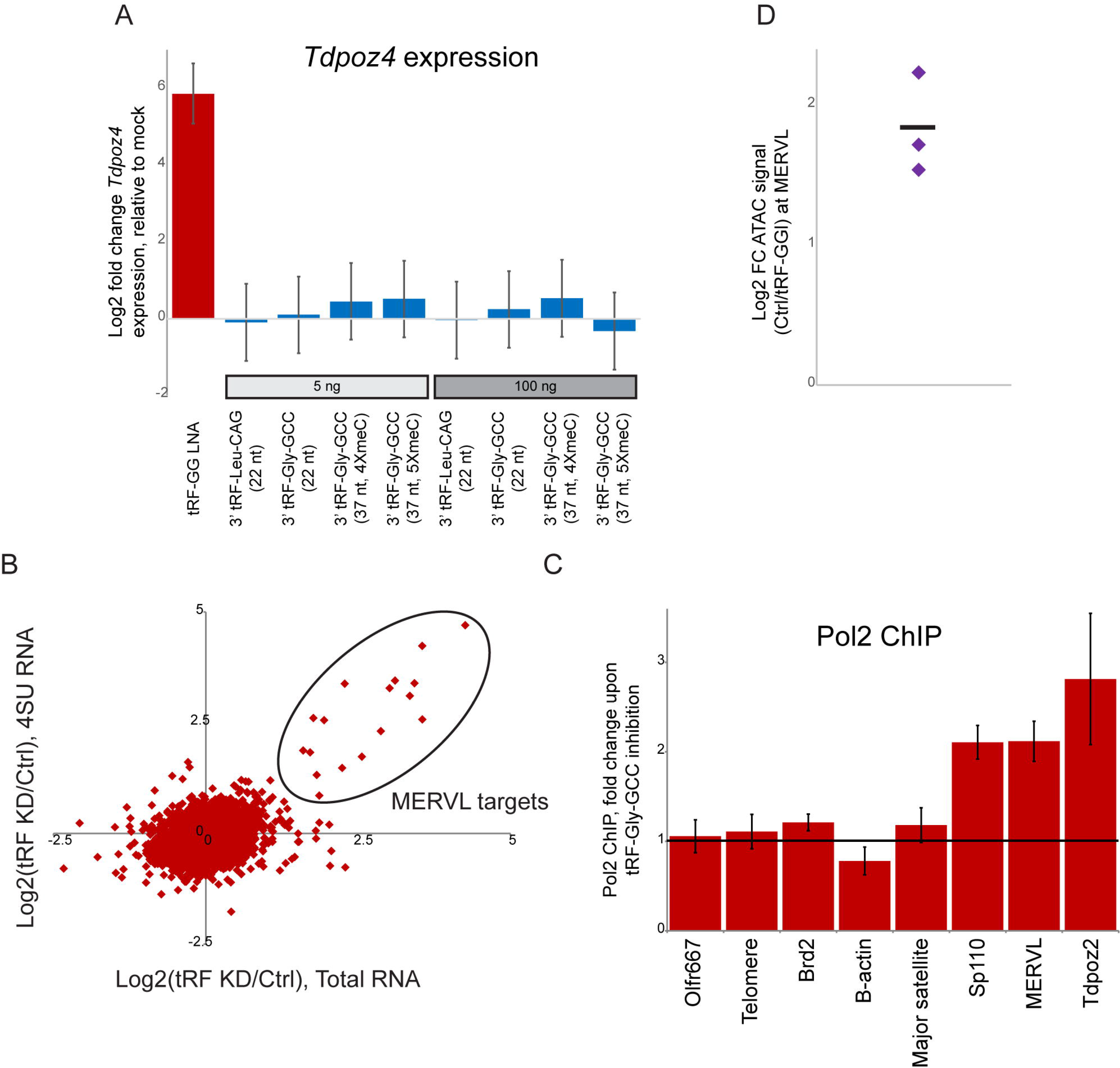
The 5’, but not the 3’, half of tRF-Gly-GCC affects MERVL target gene transcription. A) Expression of the MERVL target gene *Tdpoz4* is unaffected by several potentially relevant 3’ tRNA fragments. ES cells were transfected either with the antisense LNA targeting the 5’ tRF-Gly-GCC, or with either 5 or 100 ng of one of four synthetic 3’ tRFs. Tested oligos include two short 22 nt 3’ fragments of mature (CCA-tailed) tRNA-Gly-GCC and tRNA-Leu-CAG - the former has the potential to form a complex with 5’ tRF-Gly-GCC and so its activity could be modulated by manipulating 5’ tRF availability, while the latter was chosen due to the use of this tRNA in priming MERVL replication. We also included two longer (37 nt) 3’ tRFs derived from tRNA-Gly-GCC which differ in whether they carry 4 or 5 5meC nucleotides, as a recent study reported significant differences between these oligonucleotides in both RNA stability and structure (Zhang et al., 2018). Importantly, with the exception of the LNA targeting 5’ tRF-GG, none of the other oligos, even at elevated concentrations, had any significant effect on MERVL target gene expression in ES cell cultures. B) Metabolic labeling scatterplot showing effects of tRF-GG inhibition on total RNA (x axis) vs. newly-synthesized 4SU-labeled RNA (y axis). C) tRF-GG injection decreases chromatin accessibility at MERVL elements in preimplantation embryos. Zygotes were microinjected with a synthetic tRF-GG oligonucleotide, or control injected (H3.3-GFP mRNA), and developed to the 2-cell stage. Groups of 5 embryos were then pooled and subject to Tn5 transposition as in ATAC-Seq, then assayed by q-PCR for both MERVL and *a-tubulin*. Data show the fold decrease in accessibility for tubulin-normalized MERVL in tRF-GG-injected embryos compared to control embryos in three replicates - horizontal line shows average of the three replicates.

**Figure S2.**
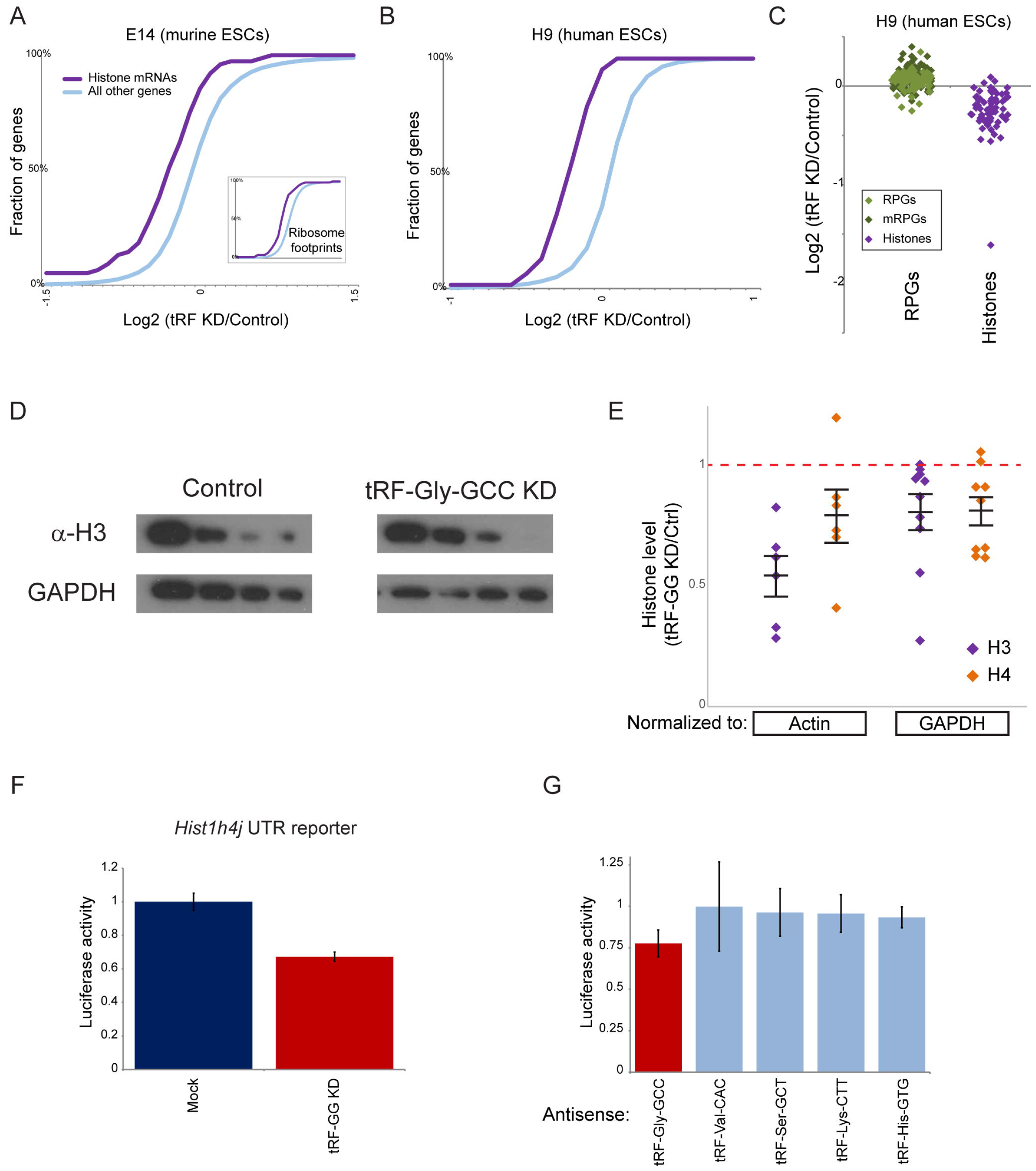
tRF KD effects on histone levels in human and mouse ESCs. A-B) Larger view of the CDF plots from **Figure 2C** for tRF-GG KD effects on histone gene expression in mouse (A) and human (B) ESCs. Inset in (A) shows effects of tRF-GG KD on ribosome occupancy (Sharma et al., 2016) of histone genes. The comparable effects on histone RNA and RBF levels suggest that tRF-GG does not directly regulate histone regulator SLBP, which regulates histone translation as well as mRNA processing/stability (Marzluff and Koreski, 2017). C) Dot plot showing effects of tRF-GG knockdown on individual histone genes (these data are replicated in **Figure 3A**). D) Example Western blots for histone H3 and GAPDH in ES cells subject to control or anti-tRF-GG transfection. The four lanes show successive 2-fold dilutions. E) Quantitation of histone levels. Dots show H3 and H4 levels for individual experiments, normalized either to β-Actin or to GAPDH, as indicated. Associated lines show mean and s.e.m. for each experiment. F) As in **Figure 2E**, for stable ES lines carrying a dual luciferase reporter with Renilla luciferase fused to the *Hist1h4j* 3’ UTR. tRF-GG inhibition resulted in a 33% decrease (p=0.000019) in luciferase expression across 8 replicates. G) Luciferase activity for the *Hist2h3b* 3’ UTR reporter, showing reporter activity in ES cells transfected with LNA antisense oligos targeting the indicated 5’ tRFs. Bars show average and standard deviation, n=4 replicates. Only tRF-GG inhibition resulted in significant (p = 0.0058) decrease in luciferase expression.

**Figure S3.**
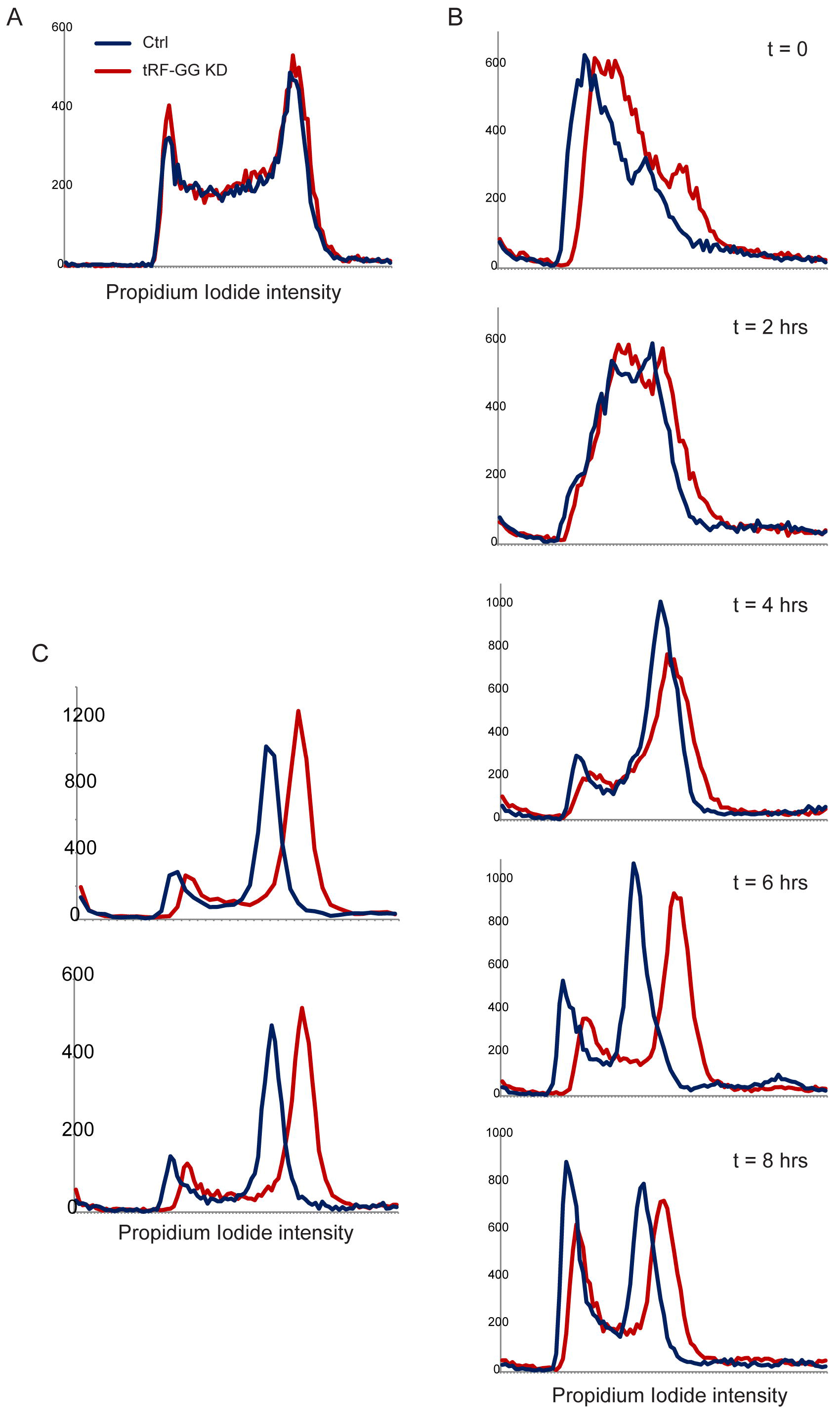
Cell cycle effects of tRF-Gly-GCC inhibition. A) Unsynchronized ES cells were either control or anti-tRF-GG LNA transfected, and 2 days later were characterized by FACS using propidium iodide staining. Nearly-identical cell cycle profiles reveal no effect of tRF-GG inhibition on the fraction of S phase cells. B) Time course of S phase progression in the presence and absence of tRF-GG inhibition. ES cells were transfected either with anti-GFP siRNAs or with the LNA antisense to tRF-Gly-GCC, then arrested in G1 phase via thymidine block for 16 hours. FACS plots show DNA content at the indicated times after release from thymidine block. tRF KD cells release into S phase and proceed to G2/M with identical kinetics, although there is a subtle G2/M exit delay in the tRF KD cells reflected in the lower frequency of G1 cells at 6 and 8 hours post-release. Note that tRF KD cells exhibit a right-shifted peak of fluorescence intensity in G2/M - this is likely due to the greater availability of naked DNA for the DNA intercalator propidium iodide. Consistent with this, reanalysis of previous FACS data for SLBP-deficient HCT cells (Jimeno-Gonzalez et al., 2015) also reveals increased fluorescence intensity in G2 cells. Taken together, our data do not support the hypothesis that increased histone levels in tRF-inhibited cells arise from an increased fraction of S phase cells. C) Additional replicates for t=6 hours after release from thymidine block.

**Figure S4.**
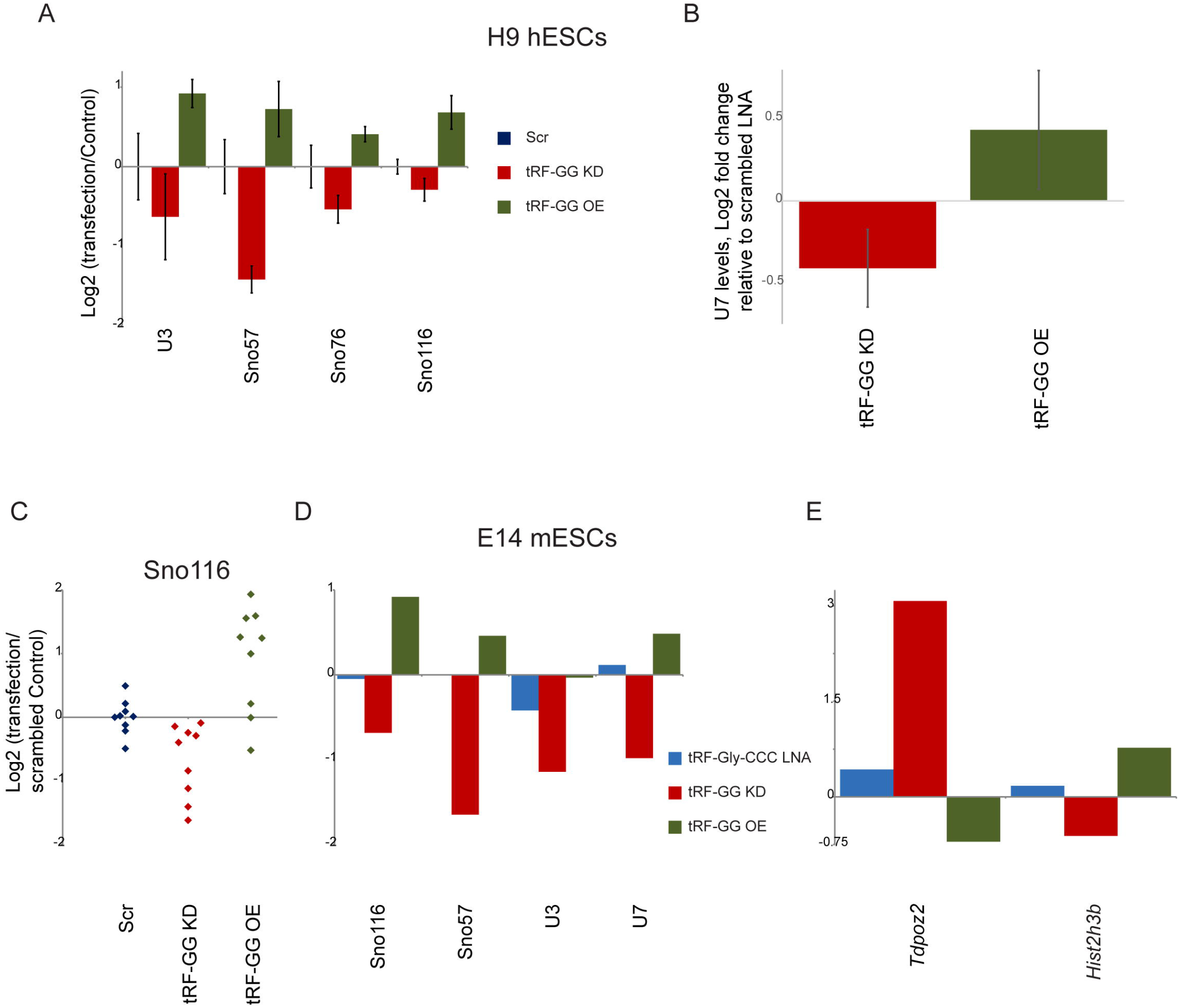
tRF-GG inhibition affects a variety of noncoding RNAs. A-B) qRT-PCR for five noncoding RNAs, normalized relative to *Actb*. Average and standard deviation are shown for three replicate experiments in H9 human ESCs, with each experiment comparing transfections with the anti-tRF-GG LNA antisense, the modified synthetic tRF-GG oligo, or a scrambled oligo control (**Methods**). Y axis shows log2 fold change relative to the scrambled oligo control. The four ncRNAs in (A) were assayed in one experiment, while the U7 q-RT-PCRs represent an independent set of transfections and so are shown separately. C-E) Data for E14 murine ESCs. (C) shows individual replicates for 9 separate experiments comparing effects of scrambled oligo, tRF-GG antisense, and tRF-GG transfection, on *Sno116* (normalized to *Actb*). (D) shows data for the four indicated noncoding RNAs (*Actb*-normalized), comparing tRF-Gly-CCC antisense, tRF-GG antisense, and tRF-GG to the scrambled oligo control. Data for Sno57, U3, and U7 were gathered in duplicates, while Sno116 data included 9 replicates for scramble, tRF-GG KD, and tRF-GG OE, and 2 replicates for tRF-Gly-CCC KD. (E) shows data for *Tdpoz2* and *Hist2h3b* in the replicate pairs shown in (D), with the overexpression of these genes in tRF-GG KD cells providing additional replication of the findings documented in **Fig.2** and previously (Sharma et al., 2016).

**Figure S5.**
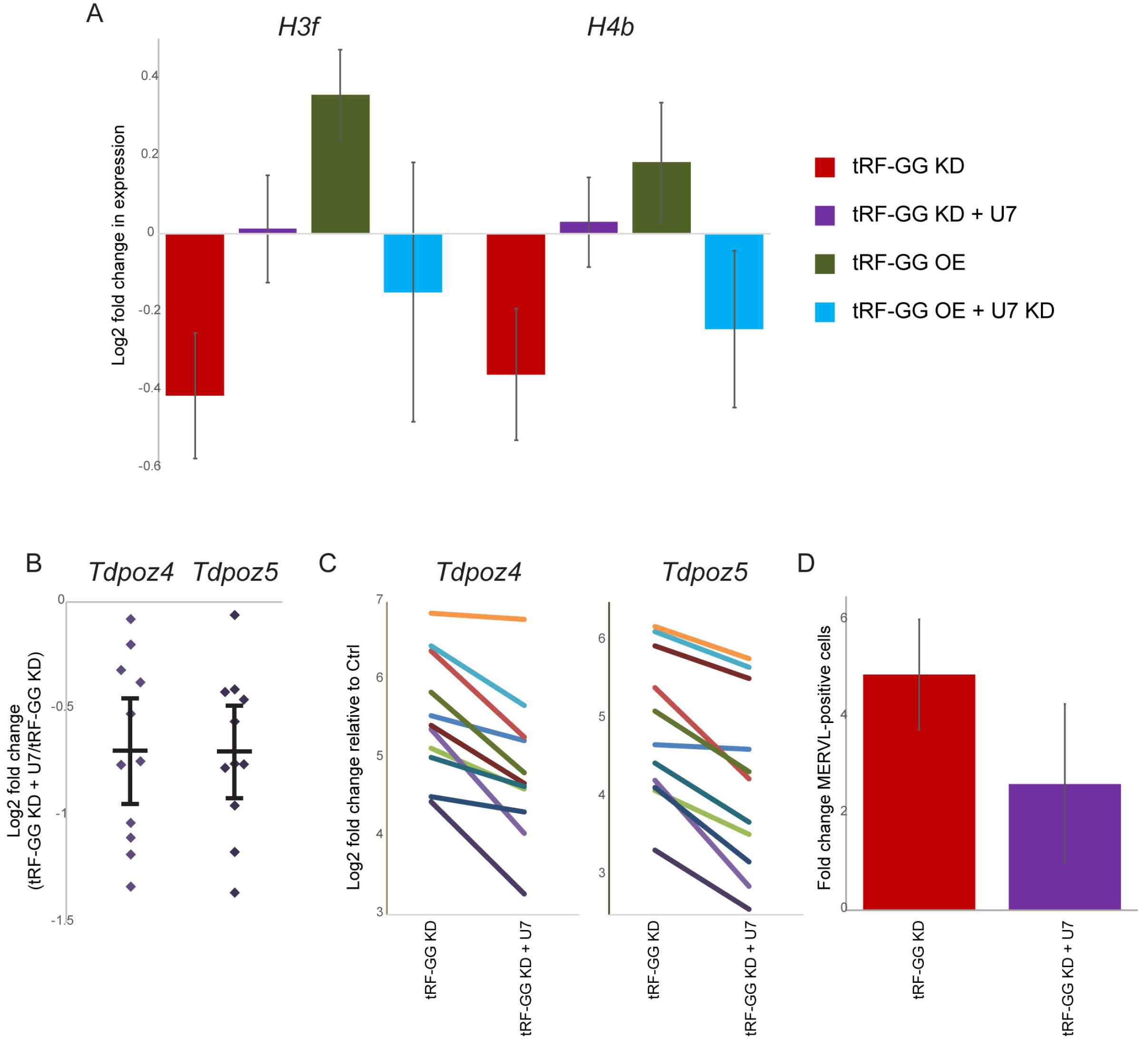
U7 suppresses effects of tRF-GG on histone mRNAs and MERVL targets. A) qRT-PCR for *HIST3F* and *HIST4B* (normalized to *ACTB*) in H9 hESCs transfected with scrambled LNA, LNA against tRF-GG with or without synthetic U7 RNA, or synthetic tRF-GG with or without antisense oligonucleotides targeting U7. Data show mean and standard deviation for three biological replicates. These data confirm the ability of tRF-GG to support histone mRNA expression, and show that the effects of tRF-GG - both positive and negative - on histone expression can be suppressed by appropriate manipulation of U7 ncRNA levels. B-C) U7 supplementation partially rescues the derepression of MERVL targets in response to tRF-GG inhibition in murine ESCs. qRT-PCR data for *Tdpoz4* and *Tdpoz5*, normalized to *Actb*, are shown for 11 biological replicates. (B) shows the relative effect of U7 on the background of tRF-GG inhibition (p=0.00015 and 4.9e-5 for *Tdpoz4* and *Tdpoz5*, respectively), with dots showing individual replicates and line+whiskers showing mean +/- s.e.m. (C) shows the effects of tRF-GG KD, with or without U7 supplementation, relative to Control ES transfections - lines connect paired transfections of the anti-tRF LNA with or without U7 supplementation. Note that y-axis does not begin at zero. In all cases, we observe a modest but consistent rescue (average log2 FC of −0.7) of *Tdpoz* repression upon U7 supplementation. The partial rescue here could result from inefficient utilization of exogenously supplied U7 ncRNA, either due to the absence of modified nucleotides in the synthetic RNA, inadequate levels of U7, or exogenous U7 not being produced at the appropriate subcellular locus (the Cajal body). D) Fold change in tdTomato-positive MERVL LTR reporter ESCs following tRF-GG knockdown with or without U7 supplementation.

**Figure S6.**
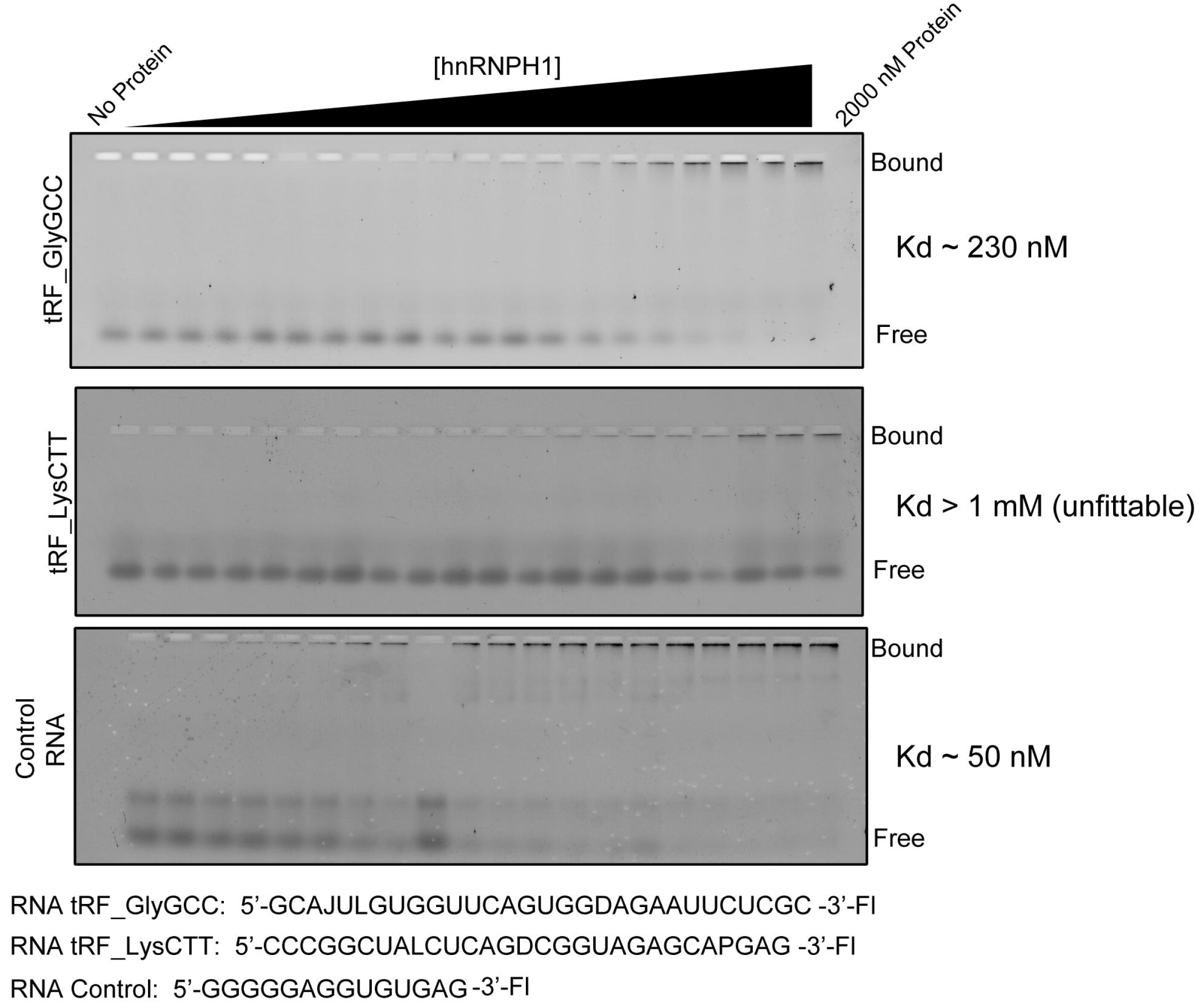
Direct binding of hnRNPH1 to tRF-Gly-GCC. Quantitative gel shift for hnRNPH1 binding to tRF-Gly-GCC is reproduced from **Figure 4E**, along with binding data for tRF-Lys-CTT and a positive control oligo (Alkan et al., 2006). Although hnRNPH1 exhibits moderately (~5-fold) weaker binding to tRF-GG than to the positive control, two considerations suggest that this binding could potentially be physiologically relevant. First, the positive control oligo was identified in a 3’ UTR shown to be regulated by hnRNPF/H. Depending on cellular growth conditions, a given tRNA fragment could easily exceed the concentration of any given 3’ UTR, so the cellular concentration of hnRNPF/H is clearly high enough for this affinity to fall into the physiologically-relevant range. Second, we note that our in vitro binding studies were performed with a synthetic 28-nt oligonucleotide bearing a FITC-modified 3’ end, while we recently found that the majority of 5’ fragments of tRNA-Gly-GCC in murine epididymis and sperm are 31-nt species bearing a cyclic 2’-3’ phosphate at the 3’ end (Sharma et al., 2018). It is thus plausible (but difficult to test given the reliance of our binding assay on 3’-modification) that hnRNPF/H binding to the naturally-produced tRF-GG is of greater affinity than that documented here.

**Figure S7.**
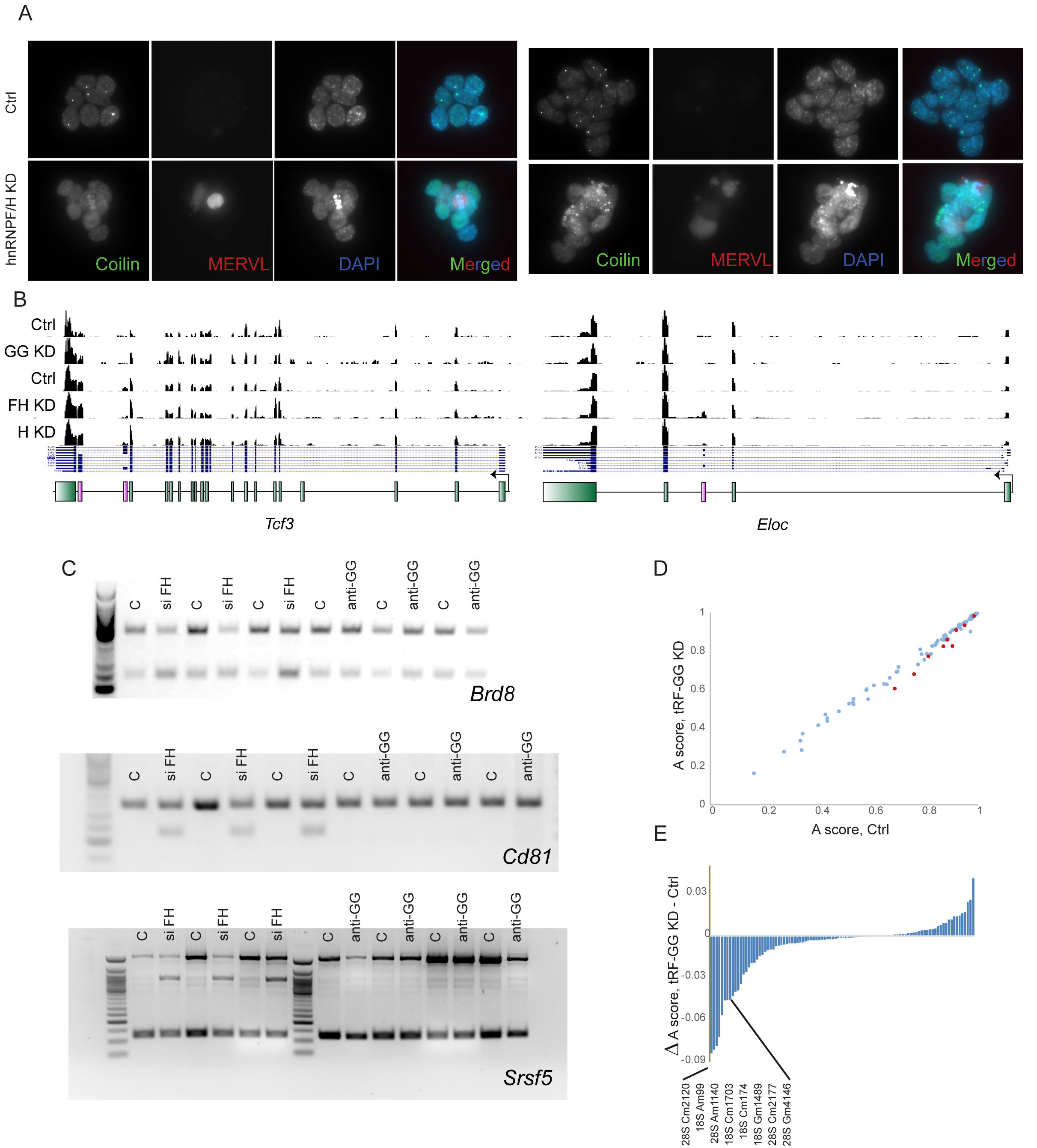
Effects of hnRNPF/H on Cajal bodies and MERVL expression. A) Immunofluorescence images of MERVL LTR:tdTomato ES cells transfected with siRNAs against either GFP or against hnRNPF and hnRNPH. Images show anti-Coilin IF, tdTomato fluorescence, DAPI counterstaining, and all three merged, as indicated. Interestingly, in preliminary immunofluorescence experiments using our MERVL LTR reporter cell line, we observed that hnRNPF/H is undetectable in spontaneously arising tdTomato-positive cells (data not shown), suggesting that hnRNPF/H degradation could potentially be involved in spontaneous ESC transitions to the 2C-like state. B) Browser shot of *Tcf3* and *Eloc* mRNA abundance in ES cells subjected to the indicated knockdowns. Alternative exon inclusion events are shown in red. C) Validation of hnRNPF/H-regulated alternative splicing events by semiquantitative PCR. Three alternative splicing events - hnRNPF/H-dependent skipping of cassette exons in *Brd8* and *Cd81*, or an hnRNPF/H-dependent detained intron in *Srsf5* - identified in our hnRNPF/H RNA-Seq data are validated here by PCR. None of these alternative splicing events was recapitulated in the tRF-GG KD samples, confirming that tRF-GG only affects a subset of hnRNPF/H functions. D-E) Altered rRNA 2’-O-methylation in tRF-GG KD cells. We carried out Ribometh-seq and calculated methylation “A scores” as previously described (Birkedal et al., 2015; Marchand et al., 2016). (D) shows a scatterplot of methylation scores for known 18S and 28S rRNA methylation sites, with significant changes in methylation highlighted in red dots. (E) shows the change in methylation level for the known rRNA nucleotides.

## REFERENCES

Alkan, S.A., Martincic, K., and Milcarek, C. (2006). The hnRNPs F and H2 bind to similar sequences to influence gene expression. The Biochemical journal 393, 361–371.

Bakowska-Zywicka, K., Kasprzyk, M., and Twardowski, T. (2016). tRNA-derived short RNAs bind to Saccharomyces cerevisiae ribosomes in a stress-dependent manner and inhibit protein synthesis in vitro. FEMS Yeast Res 16.

Birkedal, U., Christensen-Dalsgaard, M., Krogh, N., Sabarinathan, R., Gorodkin, J., and Nielsen, H. (2015). Profiling of ribose methylations in RNA by high-throughput sequencing. Angew Chem Int Ed Engl 54, 451–455.

Bogu, G.K., Vizan, P., Stanton, L.W., Beato, M., Di Croce, L., and Marti-Renom, M.A. (2015). Chromatin and RNA Maps Reveal Regulatory Long Noncoding RNAs in Mouse. Molecular and cellular biology 36, 809–819.

Buenrostro, J.D., Wu, B., Chang, H.Y., and Greenleaf, W.J. (2015). ATAC-seq: A Method for Assaying Chromatin Accessibility Genome-Wide. Curr Protoc Mol Biol 109, 21 29 21–29.

Connelly, C.M., Thomas, M., and Deiters, A. (2012). High-throughput luciferase reporter assay for small-molecule inhibitors of microRNA function. J Biomol Screen 17, 822–828.

Cossec, J.C., Theurillat, I., Chica, C., Bua Aguin, S., Gaume, X., Andrieux, A., Iturbide, A., Jouvion, G., Li, H., Bossis, G., et al. (2018). SUMO Safeguards Somatic and Pluripotent Cell Identities by Enforcing Distinct Chromatin States. Cell stem cell 23, 742–757 e748.

Couvillion, M.T., Bounova, G., Purdom, E., Speed, T.P., and Collins, K. (2012). A Tetrahymena Piwi bound to mature tRNA 3’ fragments activates the exonuclease Xrn2 for RNA processing in the nucleus. Molecular cell 48, 509–520.

Couvillion, M.T., Sachidanandam, R., and Collins, K. (2010). A growth-essential Tetrahymena Piwi protein carries tRNA fragment cargo. Genes & development 24, 2742–2747.

Deng, J., Ptashkin, R.N., Chen, Y., Cheng, Z., Liu, G., Phan, T., Deng, X., Zhou, J., Lee, I., Lee, Y.S., et al. (2015). Respiratory Syncytial Virus Utilizes a tRNA Fragment to Suppress Antiviral Responses Through a Novel Targeting Mechanism. Mol Ther 23, 1622–1629.

Dominski, Z., and Marzluff, W.F. (1999). Formation of the 3’ end of histone mRNA. Gene 239, 1–14.

Elbarbary, R.A., Takaku, H., Uchiumi, N., Tamiya, H., Abe, M., Takahashi, M., Nishida, H., and Nashimoto, M. (2009). Modulation of gene expression by human cytosolic tRNase Z(L) through 5’-half-tRNA. PloS one 4, e5908.

Gall, J.G. (2000). Cajal bodies: the first 100 years. Annu Rev Cell Dev Biol 16, 273–300.

Gebetsberger, J., Wyss, L., Mleczko, A.M., Reuther, J., and Polacek, N. (2017). A tRNA-derived fragment competes with mRNA for ribosome binding and regulates translation during stress. RNA biology 14, 1364–1373.

Gebetsberger, J., Zywicki, M., Kunzi, A., and Polacek, N. (2012). tRNA-derived fragments target the ribosome and function as regulatory non-coding RNA in Haloferax volcanii. Archaea 2012, 260909.

Goodarzi, H., Liu, X., Nguyen, H.C., Zhang, S., Fish, L., and Tavazoie, S.F. (2015). Endogenous tRNA-Derived Fragments Suppress Breast Cancer Progression via YBX1 Displacement. Cell 161, 790–802.

Guzzi, N., Ciesla, M., Ngoc, P.C.T., Lang, S., Arora, S., Dimitriou, M., Pimkova, K., Sommarin, M.N.E., Munita, R., Lubas, M., et al. (2018). Pseudouridylation of tRNA-Derived Fragments Steers Translational Control in Stem Cells. Cell 173, 1204–1216 e1226.

Heyer, E.E., Ozadam, H., Ricci, E.P., Cenik, C., and Moore, M.J. (2015). An optimized kit-free method for making strand-specific deep sequencing libraries from RNA fragments. Nucleic acids research 43, e2.

Ishiuchi, T., Enriquez-Gasca, R., Mizutani, E., Boskovic, A., Ziegler-Birling, C., Rodriguez-Terrones, D., Wakayama, T., Vaquerizas, J.M., and Torres-Padilla, M.E. (2015). Early embryonic-like cells are induced by downregulating replication-dependent chromatin assembly. Nature structural & molecular biology 22, 662–671.

Ivanov, P., Emara, M.M., Villen, J., Gygi, S.P., and Anderson, P. (2011). Angiogenin-induced tRNA fragments inhibit translation initiation. Molecular cell 43, 613–623.

Jimeno-Gonzalez, S., Payan-Bravo, L., Munoz-Cabello, A.M., Guijo, M., Gutierrez, G., Prado, F., and Reyes, J.C. (2015). Defective histone supply causes changes in RNA polymerase II elongation rate and cotranscriptional pre-mRNA splicing. Proceedings of the National Academy of Sciences of the United States of America 112, 14840–14845.

Keam, S.P., and Hutvagner, G. (2015). tRNA-Derived Fragments (tRFs): Emerging New Roles for an Ancient RNA in the Regulation of Gene Expression. Life (Basel) 5, 1638–1651.

Kim, H.K., Fuchs, G., Wang, S., Wei, W., Zhang, Y., Park, H., Roy-Chaudhuri, B., Li, P., Xu, J., Chu, K., et al. (2017). A transfer-RNA-derived small RNA regulates ribosome biogenesis. Nature 552, 57–62.

Kumar, P., Anaya, J., Mudunuri, S.B., and Dutta, A. (2014). Meta-analysis of tRNA derived RNA fragments reveals that they are evolutionarily conserved and associate with AGO proteins to recognize specific RNA targets. BMC Biol 12, 78.

Kuscu, C., Kumar, P., Kiran, M., Su, Z., Malik, A., and Dutta, A. (2018). tRNA fragments (tRFs) guide Ago to regulate gene expression post-transcriptionally in a Dicer-independent manner. RNA 24, 1093–1105.

Lee, S.R., and Collins, K. (2005). Starvation-induced cleavage of the tRNA anticodon loop in Tetrahymena thermophila. The Journal of biological chemistry 280, 42744–42749.

Lenstra, T.L., Benschop, J.J., Kim, T., Schulze, J.M., Brabers, N.A., Margaritis, T., van de Pasch, L.A., van Heesch, S.A., Brok, M.O., Groot Koerkamp, M.J., et al. (2011). The specificity and topology of chromatin interaction pathways in yeast. Molecular cell 42, 536–549.

Luo, S., He, F., Luo, J., Dou, S., Wang, Y., Guo, A., and Lu, J. (2018). Drosophila tsRNAs preferentially suppress general translation machinery via antisense pairing and participate in cellular starvation response. Nucleic acids research 46, 5250–5268.

Macfarlan, T.S., Gifford, W.D., Agarwal, S., Driscoll, S., Lettieri, K., Wang, J., Andrews, S.E., Franco, L., Rosenfeld, M.G., Ren, B., et al. (2011). Endogenous retroviruses and neighboring genes are coordinately repressed by LSD1/KDM1A. Genes & development 25, 594–607.

Macfarlan, T.S., Gifford, W.D., Driscoll, S., Lettieri, K., Rowe, H.M., Bonanomi, D., Firth, A., Singer, O., Trono, D., and Pfaff, S.L. (2012). Embryonic stem cell potency fluctuates with endogenous retrovirus activity. Nature 487, 57–63.

Machyna, M., Heyn, P., and Neugebauer, K.M. (2013). Cajal bodies: where form meets function. Wiley Interdiscip Rev RNA 4, 17–34.

Marchand, V., Blanloeil-Oillo, F., Helm, M., and Motorin, Y. (2016). Illumina-based RiboMethSeq approach for mapping of 2’-O-Me residues in RNA. Nucleic acids research 44, e135.

Marquet, R., Isel, C., Ehresmann, C., and Ehresmann, B. (1995). tRNAs as primer of reverse transcriptases. Biochimie 77, 113–124.

Martinez, G., Choudury, S.G., and Slotkin, R.K. (2017). tRNA-derived small RNAs target transposable element transcripts. Nucleic acids research 45, 5142–5152.

Marzluff, W.F., and Koreski, K.P. (2017). Birth and Death of Histone mRNAs. Trends Genet 33, 745–759.

Molla-Herman, A., Valles, A.M., Ganem-Elbaz, C., Antoniewski, C., and Huynh, J.R. (2015). tRNA processing defects induce replication stress and Chk2-dependent disruption of piRNA transcription. The EMBO journal 34, 3009–3027.

Pagano, J.M., Clingman, C.C., and Ryder, S.P. (2011). Quantitative approaches to monitor protein-nucleic acid interactions using fluorescent probes. Rna 17, 14–20.

Peng, H., Shi, J., Zhang, Y., Zhang, H., Liao, S., Li, W., Lei, L., Han, C., Ning, L., Cao, Y., et al. (2012). A novel class of tRNA-derived small RNAs extremely enriched in mature mouse sperm. Cell Res 22, 1609–1612.

Schorn, A.J., Gutbrod, M.J., LeBlanc, C., and Martienssen, R. (2017). LTR-Retrotransposon Control by tRNA-Derived Small RNAs. Cell 170, 61–71 e11.

Sharma, U., Conine, C.C., Shea, J.M., Boskovic, A., Derr, A.G., Bing, X.Y., Belleannee, C., Kucukural, A., Serra, R.W., Sun, F., et al. (2016). Biogenesis and function of tRNA fragments during sperm maturation and fertilization in mammals. Science (New York, NY 351, 391–396.

Sharma, U., Sun, F., Conine, C.C., Reichholf, B., Kukreja, S., Herzog, V.A., Ameres, S.L., and Rando, O.J. (2018). Small RNAs Are Trafficked from the Epididymis to Developing Mammalian Sperm. Developmental cell 46, 481–494 e486.

Sobala, A., and Hutvagner, G. (2013). Small RNAs derived from the 5’ end of tRNA can inhibit protein translation in human cells. RNA biology 10, 553–563.

Wang, E., Aslanzadeh, V., Papa, F., Zhu, H., de la Grange, P., and Cambi, F. (2012). Global profiling of alternative splicing events and gene expression regulated by hnRNPH/F. PloS one 7, e51266.

Wang, E., Dimova, N., and Cambi, F. (2007). PLP/DM20 ratio is regulated by hnRNPH and F and a novel G-rich enhancer in oligodendrocytes. Nucleic acids research 35, 4164–4178.

Wang, Q., and Rio, D.C. (2018). JUM is a computational method for comprehensive annotation-free analysis of alternative pre-mRNA splicing patterns. Proceedings of the National Academy of Sciences of the United States of America 115, E8181–E8190.

Wu, C.H., and Gall, J.G. (1993). U7 small nuclear RNA in C snurposomes of the Xenopus germinal vesicle. Proceedings of the National Academy of Sciences of the United States of America 90, 6257–6259.

Yamazaki, T., Liu, L., Lazarev, D., Al-Zain, A., Fomin, V., Yeung, P.L., Chambers, S.M., Lu, C.W., Studer, L., and Manley, J.L. (2018). TCF3 alternative splicing controlled by hnRNP H/F regulates E-cadherin expression and hESC pluripotency. Genes & development 32, 1161–1174.

Zhang, S., Sun, L., and Kragler, F. (2009). The phloem-delivered RNA pool contains small noncoding RNAs and interferes with translation. Plant physiology 150, 378–387.

Zhang, Y., Zhang, X., Shi, J., Tuorto, F., Li, X., Liu, Y., Liebers, R., Zhang, L., Qu, Y., Qian, J., et al. (2018). Dnmt2 mediates intergenerational transmission of paternally acquired metabolic disorders through sperm small non-coding RNAs. Nature cell biology 20, 535–540.

